# *Drosophila* kinesin-8 stabilises the kinetochore-microtubule interaction

**DOI:** 10.1101/363150

**Authors:** Tomoya Edzuka, Gohta Goshima

## Abstract

Kinesin-8 is required for proper chromosome alignment in a variety of animal and yeast cell types. However, how this conserved motor protein controls chromosome alignment remains unclear, as multiple biochemical activities, including inconsistent ones between studies, have been identified for this motor family. Here, we show that *Drosophila* kinesin-8 Klp67A possesses both microtubule (MT) plus-end-stabilising and ‐destabilising activities in addition to commonly observed MT plus-end-directed motility and tubulin-binding activity *in vitro*, and is required for stable kinetochore-MT attachment during prometaphase in S2 cells In the absence of kinesin-8^Klp67A^, abnormally-long MTs interact in an “end-on” fashion with kinetochores at normal frequency. However, the interaction was not stable and, once-attached, MTs were frequently detached. This phenotype was rescued by ectopic expression of MT plus-end-stabilising factor CLASP, but not by artificial shortening of MTs. These results suggest that MT-stabilising activity of kinesin-8^Klp67A^ is critical for stable kinetochore-MT attachment. Finally, human kinesin-8^KIF18A^ was also shown important to ensure proper MT attachment.

## Introduction

Equal segregation of sister chromatids into daughter cells relies on proper attachment of microtubules (MTs) to a specialised site on the chromosome, the kinetochore. Kinetochores consist of dozens of proteins, including those that bind to DNA or MTs, and many of them form subcomplexes for normal function (Musacchio and Desai, 2017). The Ndc80 complex is localised to the kinetochore during mitosis and functions as the major MT attachment site: “end-on” attachment of MTs to kinetochores absolutely depends on this conserved protein complex (Cheeseman et al., 2006; Musacchio and Desai, 2017; Powers et al., 2009). In yeast and animals, the Dam1 and Ska complexes, respectively, support MT binding of the Ndc80 complex (Schmidt et al., 2012; Tien et al., 2010). However, these complexes might not be the sole critical factors for MT attachment, as other MT-associated proteins, such as motor proteins, are also enriched at the kinetochore (Musacchio and Desai, 2017).

Besides attachment, kinetochores regulate the dynamics of the associated MTs. A major regulator is cytoplasmic linker associated protein (CLASP), which promotes persistent growth of kinetochore MTs (Maiato et al., 2003; Maiato et al., 2005). In its absence, MTs continuously shrink and spindles collapse (Maiato et al., 2005). *In vitro*, CLASP retards MT growth and acts as a potent inhibitor of MT “catastrophe” and as an inducer of “rescue” (Al-Bassam et al., 2010; Moriwaki and Goshima, 2016; Yu et al., 2016). Another key regulator of kinetochore MT dynamics is the kinesin-8 motor protein.

Kinesin-8 is a widely conserved kinesin subfamily. Its motor domain lies at the N-terminus, followed by coiled-coil and tail regions. The mitotic functions of kinesin-8 have been well described for budding yeast Kip3 (Cottingham and Hoyt, 1997; Straight et al., 1998; Tytell and Sorger, 2006; Wargacki et al., 2010), fission yeast Klp5/Klp6 (Garcia et al., 2001; West et al., 2002), *Drosophila* Klp67A (Gandhi et al., 2004; Goshima and Vale, 2003; Savoian et al., 2004; Savoian and Glover, 2010), and mammalian KIF18A (Mayr et al., 2007; Stumpff et al., 2008) and KIF18B (McHugh et al., 2018). Kinesin-8 is generally enriched at the outer region of the mitotic kinetochore, where plus ends of kinetochore MTs are present, and its depletion affects spindle length and chromosome alignment. In human KIF18A RNAi, the amplitude of chromosome oscillation in the abnormally-elongated spindle is dramatically elevated, such that chromosome congression cannot be achieved. In the absence of budding yeast Kip3, kinetochores are unclustered in the spindle, indicating chromosome alignment defects. Fission yeast *klp5/6* mutant also exhibits chromosome misalignment associated with Mad2-dependent mitotic delay. Overall, the loss of kinesin-8 consistently perturbs chromosome alignment in a variety of cell types.

Despite the conserved phenotype and localisation associated with kinesin-8, its biochemical activity towards MTs is inconsistent between reports. The best-studied budding yeast Kip3 has plus-end-directed, processive motility and also has strong MT depolymerising activity; it can depolymerise MTs stabilised by GMPCPP (non-hydrolysable GTP) and promote catastrophe (growth-to-shrinkage transition) in dynamic MTs (Gupta et al., 2006; Varga et al., 2006). The C-terminal tail has MT‐ and tubulin-binding activities, which allow this motor to crosslink and slide antiparallel MTs (Su et al., 2013; Su et al., 2011). However, MT depolymerisation activity has not been detected for fission yeast proteins Klp5/Klp6 and MT nucleation activity has been reported instead (Erent et al., 2012). Humans have two mitotic kinesin-8s, KIF18A and KIF18B, and kinetochore function has been observed for KIF18A. KIF18A, like Kip3, exhibits processive motility towards plus ends, and accumulates at plus ends on its own (Du et al., 2010; Mayr et al., 2007). The tail region of KIF18A has MT and tubulin affinity, which is similar to Kip3 (Mayr et al., 2011; Weaver et al., 2011). However, its impact on MT dynamics has been controversial. In one study, KIF18A was concluded to have MT depolymerising activity, based on its depolymerisation activity towards stabilised MTs (Mayr et al., 2007). In another study, however, this activity was reported to be undetectable, and instead, it dampened MT dynamicity; KIF18A suppressed both growth and shrinkage of MTs (Du et al., 2010). Although the former activity is more consistent with Kip3, the latter activity appears to be more congruous with the cellular phenotype associated with KIF18A (Stumpff et al., 2008).

In the present study, we investigated Klp67A, the sole mitotic kinesin-8 in *Drosophila*. In addition to conserved MT-based motility, we identified both MT stabilising and destabilising activities in a single experimental condition. Functional analysis in the S2 cell line indicated that, with these two activities, kinesin-8^Klp67A^ not only regulates MT length, but also stabilises kinetochore-MT attachment.

## Results

### Kinesin-8^Klp67A^ shows plus-end-directed motility and tubulin-binding activity *in vitro*

Generally observed biochemical activities among mitotic kinesin-8 motors are processive motility (Gupta et al., 2006; McHugh et al., 2018; Stumpff et al., 2011; Varga et al., 2006) and tubulin binding at the non-motor region (Mayr et al., 2011; Su et al., 2011; Weaver et al., 2011). To determine the biochemical activity of *Drosophila* kinesin-8^Klp67A^, we purified recombinant GFP-tagged full-length protein (Fig. S1A). First, we performed single motor motility assays and found that kinesin-8^Klp67A^-GFP is a processive motor with mean velocity 25 ± 0.7 µm/min (± SEM) (Fig. 1A, B, Movie 1). The GFP signal quickly accumulated at one end of MTs: therefore, to visualize how each motor accumulates at the MT end, we photo-bleached the pre-accumulated kinesin-8^Klp67A^-GFP signals at the end, followed by observing unbleached motors. In 32 out of 33 cases, we observed that the motile motor on the MT lattice reached and stayed at the end of the MT polymers (Fig. 1A, Movie 1). Thus, kinesin-8^Klp67A^-GFP processively moved to the end and resided there for a certain period; this behaviour is identical to that observed for Kip3 (Varga et al., 2009). Next, we determined the directionality of motility by localising kinesin-8^Klp67A^-GFP on MTs that underwent gliding by the plus-end-directed kinesin-1 motor (Fig. 1C). Kinesin-8^Klp67A^-GFP was enriched at the trailing end of MTs, indicating that kinesin-8^Klp67A^ is a plus-end-directed motor, as seen with other kinesin-8s (Fig. 1D).

**Figure 1.**
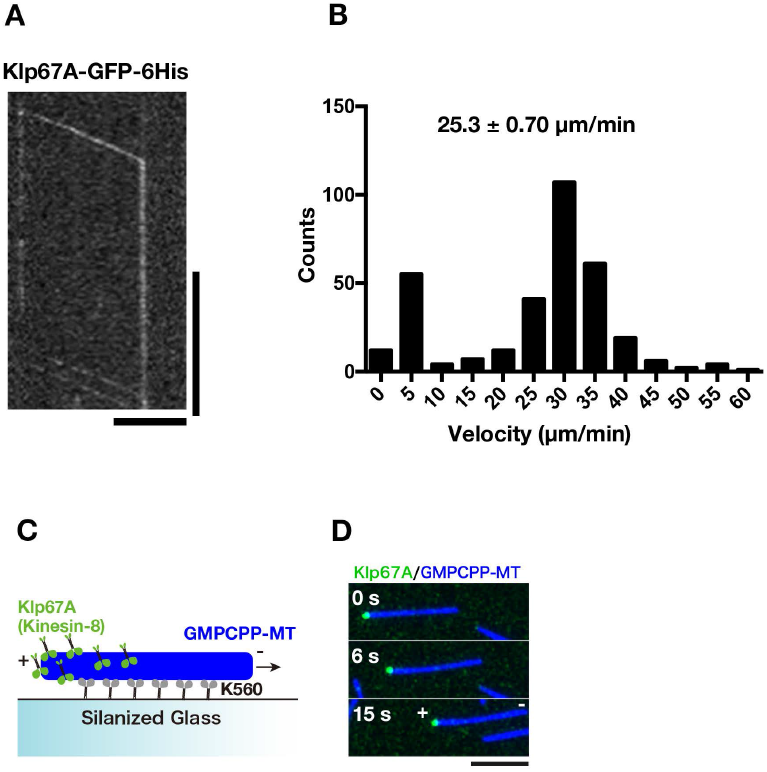
Kinesin-8^Klp67A^ is a processive plus-end-directed motor. **(A)** Kymograph showing processive motility of kinesin-8^Klp67A^-GFP (100 pM) along MTs and accumulation at the MT end. Kinesin-8^Klp67A^-GFP enriched at the MT end was first photo-bleached, followed by observation of unbleached kinesin-8^Klp67A^-GFP. Horizontal bar, 5 μm; Vertical bar, 20 s. **(B)** Plot of kinesin-8^Klp67A^-GFP run velocity, calculated based on kymographs (n = 331). Immobile GFP signals were not counted (the “0” column represents velocity between 0 and 5 μm/min). **(C, D)** Gliding of stabilised MTs (blue) by the plus-end-directed human kinesin-1 motor (K560 construct; non-fluorescent). Rightward motility of MTs indicate that the left end corresponds to MT plus end. Kinesin-8^Klp67A^-GFP (green, 4 nM) accumulated specifically at the plus end.

To test if kinesin-8^Klp67^ binds to tubulin, like Kip3 and KIF18A, we performed sucrose gradient centrifugation of recombinant kinesin-8^Klp67A^-GFP in the presence and absence of tubulin. Kinesin-8^Klp67A^-GFP was co-fractionated with tubulin as a larger complex, indicating that kinesin-8^Klp67A^ directly binds to tubulin (Fig. 2A). Next, to verify this finding and further identify the region responsible for tubulin binding, we performed tubulin recruitment assay, in which kinesin-8^Klp67A^ (full-length and truncations) was localised along the stabilised MT seed and fluorescently-labelled free tubulin was added and observed (Fig. 2B, C, S1B). Consistent with the sucrose gradient centrifugation analysis, tubulin was efficiently recruited to the MT seed by kinesin-8^Klp67A^ full-length (Fig. 2D, E). C-terminally truncated “tail-less” construct (1–612 a.a.), which was shown to rescue the spindle length and chromosome alignment phenotypes (Savoian and Glover, 2010), also recruited tubulin onto MT seeds, albeit less efficiently than full-length. However, when non-motor regions were entirely eliminated, tubulin was hardly recruited. These results indicate that kinesin-8^Klp67A^ binds to tubulin at the non-motor region and suggest that the binding is important for kinesin-8^Klp67A^ function.

**Figure 2.**
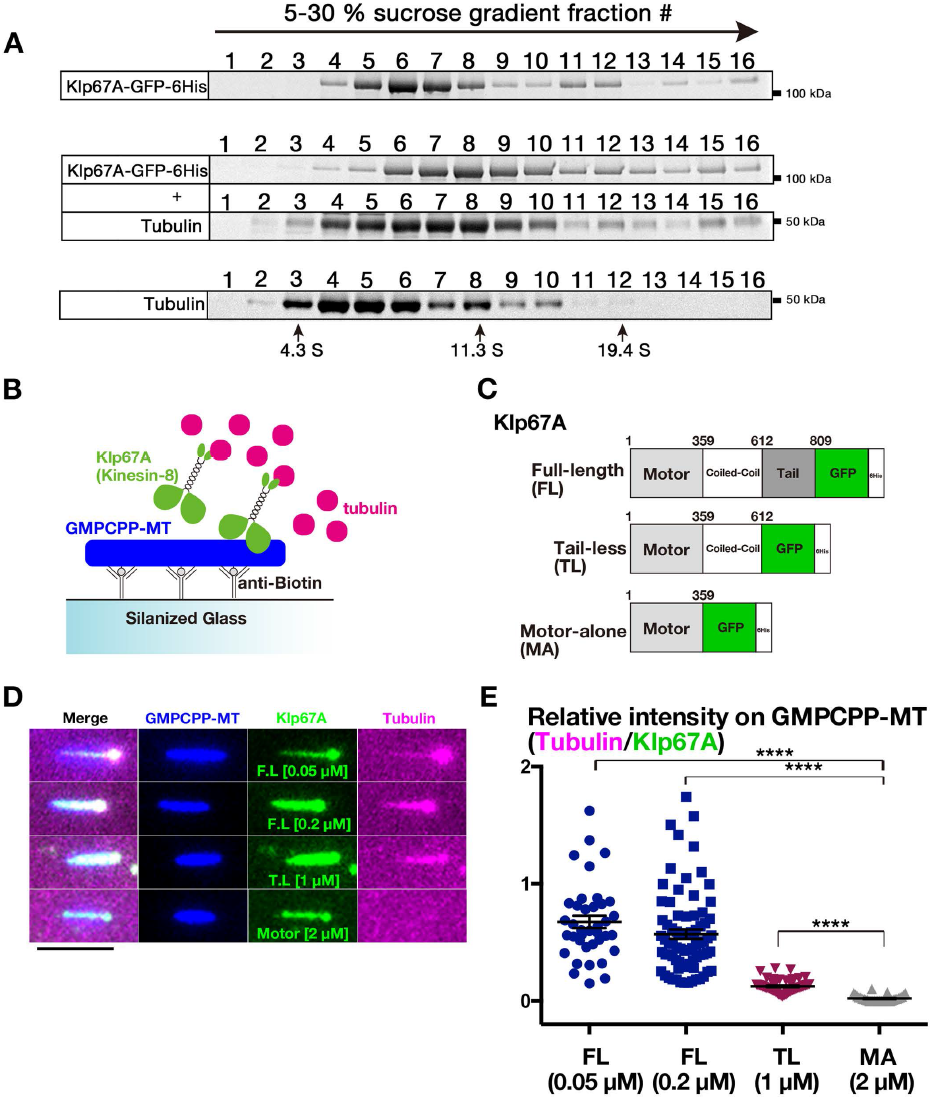
Tubulin-binding activity of kinesin-8^Klp67A^. **(A)** Co-fractionation of kinesin-8^Klp67A^-GFP and tubulin after sucrose gradient centrifugation. Each fraction was subjected to SDS-PAGE, followed by staining with Sypro Ruby. **(B-D)** Tubulin recruitment by kinesin-8^Klp67A^. Tubulin (10 μM, magenta) and kinesin-8^Klp67A^ (green; full-length, tail-less [1–612 a.a.], motor-alone [1–359 a.a.]), which bound to GMPCPP-stabilised MTs (blue), were mixed, and tubulin localisation along MTs was investigated. Bar, 5 μm. **(E)** Quantification of tubulin intensity on the MT seed. Each dot represents a value obtained from a single MT and error bars represent SEM. FL vs. MA: p = 3.8 x10^-14^, TL vs. MA: p = 3.8 × 10^-10^ by Games-Howell test, n = 38 (50 nM) + 77 (200 nM) (full-length), 33 (motor-alone), 58 (tail-less).

### Kinesin-8^Klp67A^ has both MT-destabilising and ‐stabilising activities *in vitro*

We next determined the effect of kinesin-8^Klp67A^ on MT polymerisation dynamics in an *in vitro* assay (Fig. 3A). When we mixed 5, 10, 20, or 50 nM kinesin-8^Klp67A^ with GMPCPP-stabilised MT seeds and free tubulin (10 µM), dynamic MTs from the seeds were rarely observed or non-existent at 20 nM or 50 nM, respectively (Fig. 3B). Furthermore, at high concentrations, the excess force exerted by kinesin-8^Klp67A^ glided and bundled MT seeds. In the subsequent experiment, we used 5 or 10 nM kinesin-8^Klp67A^.

**Figure 3.**
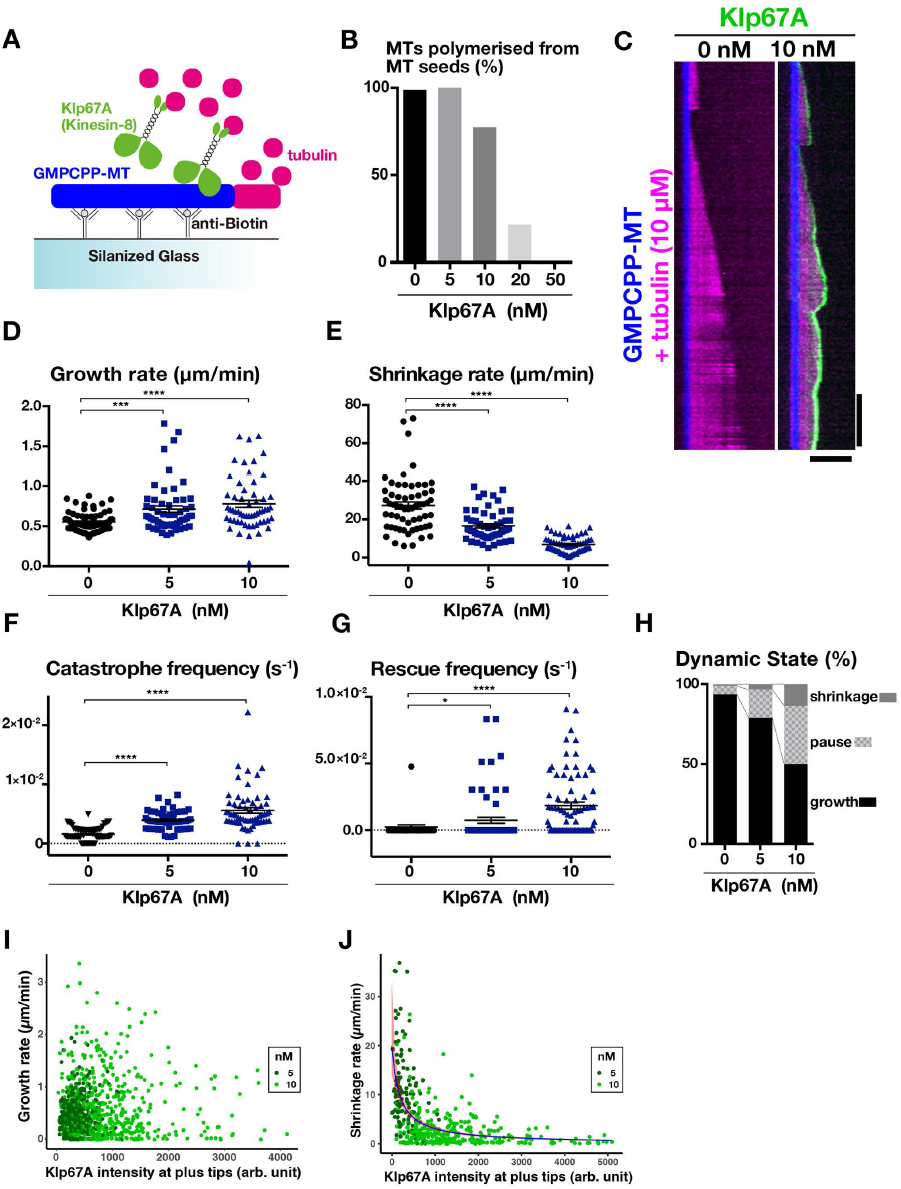
Kinesin-8^Klp67A^ regulates MT plus-end dynamics in vitro. **(A)** Schematic presentation of the in vitro MT polymerisation assay. **(B)** Inhibitory effect of kinesin-8^Klp67A^ on MT polymerisation from the seed (n = 84, 74, 84, 74, 94 from left to right). **(C)** MT dynamics are represented by kymographs. MTs stabilised with GMPCPP are coloured blue, dynamics MTs are magenta, and kinesin-8^Klp67A^-GFP (0 or 10 nM) is green. Horizontal bar, 5 μm; Vertical bar, 120 s. **(D-H)** Parameters of MT plus end dynamics. Experiments were performed twice and the combined data are displayed (the change in rate/frequency by kinesin-8^Klp67A^ was reproduced). N = 41 + 33 (0 nM), 25 + 31 (5 nM), 20 + 41 (10 nM). Each dot represents a value obtained from a single MT and error bars represent SEM. Note that rescue was rarely observed in the absence of kinesin-8^Klp67A^. p < 0.0001 (****), p < 0.002 (***), or p < 0.03 (*) by Games-Howell **(D-F)** or Steel Dwass (G) tests. (I, J) Correlation between the amount of kinesin-8^Klp67A-GFP^ at the tip and growth (I; n = 397 [5 nM] and 494 [10 nM]) or shrinkage (J; n = 116 [5 nM] and 293 [10 nM]) rate. Negative correlation was found for shrinkage rate (p < 2 x 10^-16^, likelihood ratio test).

First, we could not observe any MT depolymerisation activity of kinesin-8^Klp67A^ towards GMPCPP-stabilised MTs, which differs from that for yeast Kip3 (Gupta et al., 2006; Varga et al., 2006), human KIF18A reported by (Locke et al., 2017; Mayr et al., 2007), or the bona fide MT-depolymerising *Drosophila* kinesin-13^Klp10A^ (Rogers et al., 2004) that was used as a positive control in our experiments (Fig. S1C). However, we could not exclude the possibility that kinesin-8^Klp67A^ has a weak seed depolymerisation activity that was undetectable with this motor concentration.

Next, we quantified the dynamics parameters in the presence of 5 or 10 nM kinesin-8^Klp67A^ (Fig. 3C–G). As expected from the spindle lengthening phenotype, kinesin-8^Klp67A^ elevated the catastrophe frequency of the MTs. Interestingly, kinesin-8^Klp67A^ also increased the rescue frequency and slowed down shrinkage under the same assay conditions, which resulted in more frequent MT pausing (Fig. 3H). Thus, we observed both MT-stabilising and ‐destabilising effects reported for kinesin-8^KIF18A^ by (Du et al., 2010) and (Locke et al., 2017; Mayr et al., 2007), respectively. The growth rate was increased in a slight but statistically significant manner, which is consistent with a recent report concerning human kinesin-8^KIF18B^ (McHugh et al., 2018).

Finally, we analysed the correlation between the amount of kinesin-8^Klp67A^-GFP at the plus end and the MT growth/shrinkage rate. We measured and plotted GFP intensity at the tip and the velocity of the MTs (Fig. 3I, J). GFP intensity at the shrinking tip was, on average, slightly higher than that at the growing tip (Fig. S1D). Whether this small difference is of physiological relevance is unclear. Interestingly, there was a strong anti-correlation between GFP intensity and shrinking velocity (p < 2 × 10^-16^, gamma regression, likelihood-ratio tests; Fig. 3J), whereas no clear correlation was identified for growth (p = 0.99; Fig. 3I). The data is consistent with the model that kinesin-8^Klp67A^ accumulated at the tip induces catastrophe but simultaneously prohibits drastic shrinkage. On the other hand, the mechanism by which kinesin-8^Klp67A^ increases the MT growth rate is unclear.

### Kinesin-8^Klp67A^ depletion causes instability of kinetochore-MT attachment

To explore the function of kinesin-8^Klp67A^ in the spindle, we observed its RNAi phenotype in living S2 cells (RNAi knockdown was confirmed by immunoblotting; Fig. S2A). In addition to chromosomes and MTs, we traced GFP-Rod: GFP-Rod accumulates at unattached or laterally-attached kinetochores and, once MTs are attached in an end-on fashion, it is transported away from the kinetochore along the MTs (Basto et al., 2004; Gluszek et al., 2015). Therefore, GFP-Rod exhibits “streaming” upon MT end-on attachment, while residual proteins are still visible at the kinetochore; thus, it serves as an ideal marker for kinetochore dynamics as well as for its attachment status (Fig. 4A). To precisely monitor and evaluate the dynamics of individual chromosome and GFP-Rod signals in the uniformly-shaped spindle, we induced monopolar spindles by depleting Klp61F, the kinesin-5 motor protein required for spindle bipolarisation. MTs were stained with the SiR-tubulin dye. In control cells that were singly depleted of kinesin-5^Klp61F^, we observed that the majority of sister kinetochores were attached to MTs from the pole (“syntelic” attachment), with weak GFP-Rod signals detected (Fig. 4B, Movie 2). The syntelically attached kinetochores were static and scarcely changed position during the observation time. In some instances, we observed “monotelic” attachment at the beginning, where one of the sister kinetochores was not associated with MTs and, therefore, a strong GFP-Rod signal was detected (Fig. 4C, 12 s). However, they were converted into syntelic attachments during imaging, which was characterised by GFP-Rod streaming along the newly formed kinetochore MTs (Fig. 4C, 153 s). Once syntelic attachment was established, MTs were rarely dissociated from kinetochores (Fig. 4D, control).

**Figure 4.**
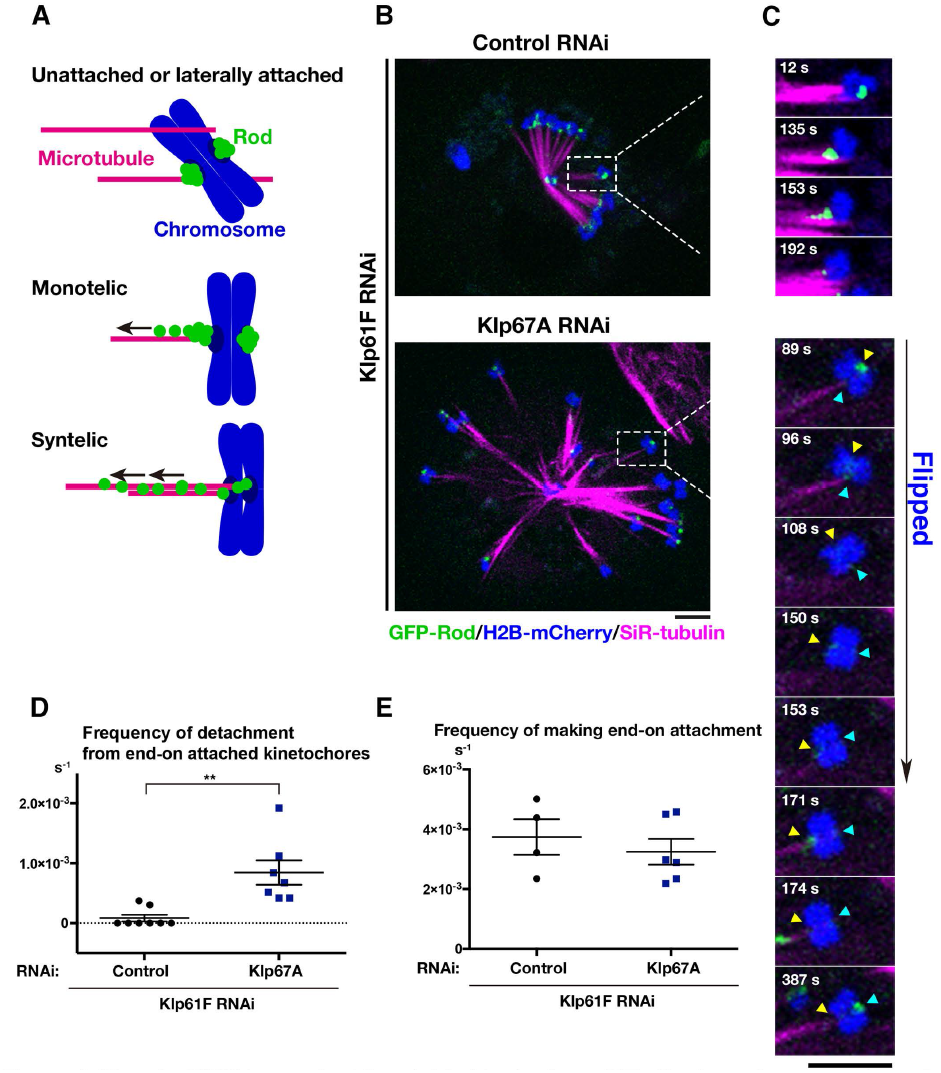
Kinesin-8^Klp67A^ is required for stable kinetochore-MT attachment: monopolar spindle. **(A)** Diagram of GFP-Rod and MT attachment mode. **(B)** Monopolar spindles formed after RNAi of kinesin-5^Klp61F^ (top) or double kinesin-5^Klp61F^/ kinesin-8^Klp67A^ (bottom). Blue, chromosome (H2B-mCherry); Green, GFP-Rod; Magenta, MT (SiR-tubulin). **(C)** (Top) Monotelic-to-syntelic conversion in the control monopolar spindle; upon new MT association, GFP-Rod stream was observed along the kinetochore MT (153 s), and subsequently, the GFP signal diminished at the kinetochore (192 s). (Bottom) Chromosome flipping was observed in the absence of kinesin-8^Klp67A^(89-171 s; sister kinetochores are indicated by yellow and blue arrows). Monotelic-to-syntelic conversion was also observed for this chromosome (96 s). **(D)** Increased frequency (event # per chromosome per sec) of MT detachment in the absence of kinesin-8^Klp67A^ (p < 0.009, Welch’s t-test). Events were counted when MTs terminated end-on attachment. RNAi and imaging were performed four times, data were quantitatively analysed twice, and the two datasets were combined. Each dot in the graph represents mean frequency for a cell. A total of 86 chromosomes in 8 cells (control) and 56 chromosomes in 8 cells (kinesin-8^Klp67A^ RNAi) were analysed. **(E)** Frequency (event # per chromosome per sec) of newly acquired end-on attachment that was indicated by GFP-Rod stream along kinetochore MTs. The attachment number was divided by the total time the chromosomes spent in a monotelic or unattached state. Each dot in the graph represents mean frequency for a cell. A total of 18 chromosomes in 4 cells (control) and 18 chromosomes in 6 cells (kinesin-8^Klp67A^ RNAi) were analysed. Error bars indicate SEM. Bars, 5 μm.

When kinesin-5^Klp61F^ and kinesin-8^Klp67A^ were co-depleted, monopolar spindles with much longer MTs were assembled (Fig. 4B). In addition, kinetochore dynamics were dramatically different in these cells (Movie 2). Some kinetochores were visibly motile, and monotelic attachment was more frequently observed for those chromosomes. Most of the unattached kinetochores acquired end-on attachment during the imaging period, as indicated by GFP-Rod streaming (e.g. Fig. 4C, 171–174 s), although the associations were often transient. Quantification indicated detachment of MTs from the kinetochore was significantly more frequently observed in the absence of kinesin-8^Klp67A^, whereas end-on attachment event was detected at a frequency similar to that in control cells (Fig. 4D, E). Interestingly, MT detachment was usually associated with chromosome flipping, where a sister kinetochore originally distal from the pole was flipped to face the pole (e.g. Fig. 4C, 89–153 s).

To verify that the observed phenotype is not an artefact of SiR-tubulin staining, we observed a cell line that expressed GFP-Rod, H2B-mCherry, and EB1-SNAP (fluorescent SiR-SNAP was added to the medium), in which EB1 served as a marker of kinetochore MTs (Fig. S2B). In addition, we simply observed GFP-Rod and H2B-mCherry without MT markers (Fig. S2D). In the absence of kinesin-8^Klp67A^, chromosome flipping was 19-fold more frequently observed than in control cells (n = 15 and 27), confirming the role of kinesin-8^Klp67A^ in stable kinetochore-MT attachment in the monopolar spindle (Fig. S2C).

We next observed GFP-Rod behaviour in the bipolar spindle with and without kinesin-8^Klp67A^ (Fig. 5). As previously reported, abnormally-elongated spindles with unaligned chromosomes were observed in the absence of kinesin-8^Klp67A^. Since MTs were crowded in the spindle, it was impossible to observe MT attachment status for most kinetochores. Nevertheless, when we focused on completely unaligned chromosomes that were remote from the main body of the spindle, we observed a phenotype similar to that observed in the monopolar assay. In the control cell displayed in Fig. 5A and Movie 3, a chromosome (arrow) was not immediately captured by MTs and remained near the pole; it had strong GFP-Rod signals. However, MTs were generated independent of centrosomes and bound to the kinetochore (63 s). Once those MTs were formed, the flow of GFP-Rod was visible, concomitant with the decrease of the kinetochore signal intensity (93–180 s). Thus, a bi-oriented chromosome with “amphitelic” attachment was finally observed, and it was translocated towards the spindle equator (375 s). In contrast, MT attachment was unstable and MT detachment was observed in kinesin-8^Klp67A^-depleted cells. In the case displayed in Fig. 5B and Movie 3, a misaligned chromosome (arrow) achieved amphitelic attachment at 114 s, as evident by GFP-Rod stream along chromosome-bound MTs, but then such a configuration was disrupted at 198 s and the chromosome was flipped (198–285 s). MT detachment frequency was quantified in Fig. 5C.

**Figure 5.**
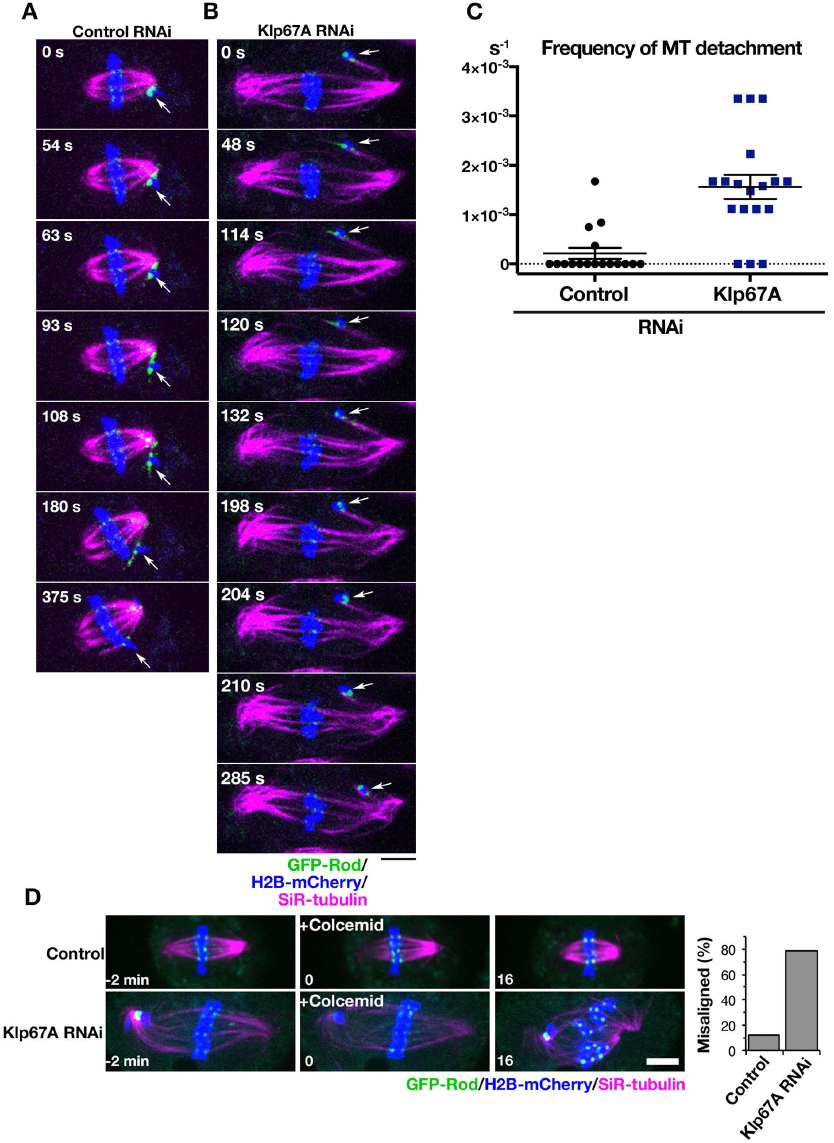
Kinesin-8^Klp67A^ is required for stable kinetochore-MT attachment: bipolar spindle. **(A, B)** Time-lapse imaging of spindle MTs (magenta; SiR-tubulin staining), chromosomes (blue; H2B-mCherry), and GFP-Rod (green) after control or kinesin-8^Klp67A^ RNAi. The behaviour of an initially unaligned chromosome (arrow) was dramatically different. **(C)** Increased frequency (event # per cell per sec) of MT detachment in the absence of kinesin-8^Klp67A^ (p < 0.001, Welch’s t-test). Events were counted when MTs terminated end-on attachment. RNAi and imaging were performed 3 times, data were quantitatively analysed twice, and the two datasets were combined. Error bars represent SEM. (D) Sensitivity to MT depolymerisation. One or more chromosomes at the metaphase plate became unaligned upon colcemid treatment at much higher frequency in the absence of kinesin-8^Klp67A^. Bars, 5 μm.

Finally, we treated the metaphase cells with a MT destabilising drug and preferentially depolymerised non-kinetochore MTs. In control cells, kinetochore MTs kept the chromosomes aligned at the metaphase plate in 88% of the cases (n = 25) for ≥ 15 min, as expected (Goshima et al., 2008). In contrast, at least one chromosome that had been located at the metaphase plate was misaligned in 78% of the cells in the absence of kinesin-8^Klp67A^ (Fig. 5D).

From these results, we concluded that the kinetochore-MT association becomes unstable in the absence of kinesin-8^Klp67A^.

### Artificial MT destabilisation does not rescue the kinesin-8^Klp67A^-depleted phenotype

Attachment instability might be the consequence of longer MTs in the absence of kinesin-8^Klp67A^. To test this possibility, we depleted kinesin-5^Klp61F^ and Dgt6, an essential subunit of the MT amplifier augmin, to generate monopolar spindles with long MTs; MTs are elongated in this condition due to the reduction of MT nucleation sites within the spindle (Goshima et al., 2008). Augmin^Dgt6^/kinesin-5^Klp61F^ RNAi-treated cells indeed exhibited much longer and pendulous MTs, similar to those observed after kinesin-8^Klp67A^ depletion (Fig. S3A, Movie 4). Monotelic and syntelic attachments were both observed, similar to that seen in kinesin-8^Klp67A^-depleted cells. However, once-attached, MTs were more persistent and chromosome flipping was rarely observed (Fig. 6D).

**Figure 6.**
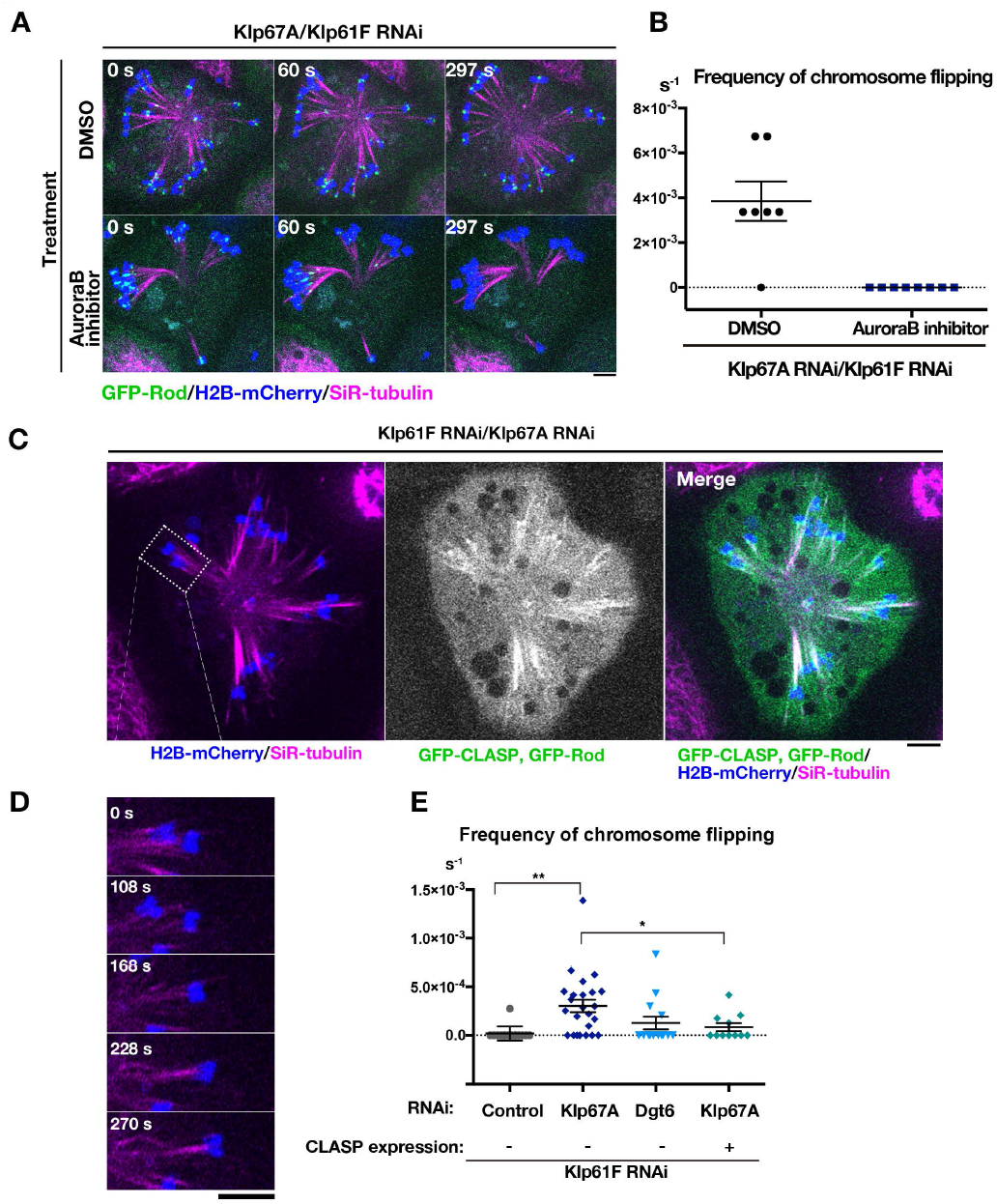
MT attachment stability was restored by inhibiting Aurora B kinase or CLASP^Mast/Orbit^ overexpression. **(A)** Monopolar spindles were induced after double kinesin-5^Klp61F^/kinesin-8^Klp67A^ RNAi and treated with 20 μM Binucleine-2, an inhibitor of *Drosophila* Aurora B kinase, or control DMSO. (B) Frequency of chromosome flipping (per cell, per sec, ±SEM) in Binucleine-2 treated cells (n = 7 [control DMSO-treated] and 8 cells [Binucleine-2-treated]). (C) GFP-CLASP^Mast/Orbit^ overexpression in the monopolar spindle depleted of kinesin-5^Klp61F^ and kinesin-8^Klp87A^. (D) Two representative chromosomes (inset in C), Chromosomes were static when GFP-CLASP^Mast/Orbit^ was overexpressed. (E) Frequency of chromosome flipping (per chromosome, per sec, ±SEM). Expression of GFP-CLASP^MESt/Orbit^ significantly reduced the frequency of flipped chromosomes (**p < 0.0013, *p < 0.034, Games-Howell test). RNAi (and overexpression) were performed two or more times, and combined data are presented. Each dot in the graph represents mean frequency for a cell. A total of 168 chromosomes in 15 cells (control), 364 chromosomes in 24 cells (kinesin-8^Klp67A^ RNAi), 128 chromosomes in 13 cells (augmin^Dgt6^ RNAi), 128 chromosomes in 11 cells (kinesin-8^Klp67A^ RNAi and GFP-CLASP^Mast/Orbit^ overexpression) were analysed. Bars, 5 μm.

To further exclude the possibility that abnormally-elongated MTs due to reduced catastrophe are the major cause of MT attachment instability, we exposed a low dosage of colcemid to kinesin-8^Klp67A^ RNAi-treated cells and shortened the bipolar spindle lengths to the control levels (Fig. S3B, C). Unaligned chromosomes were still frequently observed and mitosis was significantly delayed (Fig. S3B, D). These results suggested that regulation of MT length alone cannot explain the attachment instability of kinesin-8^Klp67A^-depleted cells.

### Aurora B kinase inhibition or CLASP^Mast/Orbit^ overexpression rescues attachment instability caused by kinesin-8^Klp67A^ depletion

In mammalian cells, inhibition of Aurora B kinase stabilises syntelic attachment, at least partly due to dephosphorylation of Ndc80: Ndc80 is a critical kinetochore component for MT binding and its MT binding affinity is decreased by Aurora B phosphorylation (Cheeseman et al., 2002; DeLuca et al., 2006; Lampson and Grishchuk, 2017; Musacchio and Desai, 2017). We added the inhibitor of *Drosophila* Aurora-B, Binucleine-2 (Smurnyy et al., 2010), to S2 cells depleted of kinesin-8^Klp67A^/kinesin-5^Klp61F^, and observed that most kinetochores stably attached MTs in a syntelic manner, as indicated by a decrease in GFP-Rod signals (Fig. 6A, B, Movie 5). The result indicates that kinetochores retain an ability to bind stably to MTs in the absence of kinesin-8^Klp67A^, when Aurora B activity is low. Our interpretation is that the MT-binding potential of the dephosphorylated form of Ndc80 complexes is preserved in the absence of kinesin-8^Klp67A^.

In the absence of kinesin-8^Klp67A^, persistent poleward motility of chromosomes was observed, which would involve kinetochore MT shrinkage (Movie 2). We hypothesised that the MT stabilisation activity of kinesin-8^Klp67A^, namely slowing down MT shrinkage and inducing rescue/pausing, is critical for MT attachment stability. If that were the case, we reasoned that MT stabilisation by other means might partially suppress the MT detachment phenotype. To this end, we expressed *Drosophila* CLASP (also called Mast or Orbit) in kinesin-8^Klp67A^/kinesin-5^Klp61F^ RNAi cells and observed the consequent chromosome dynamics. Since we attached GFP to CLASP^Mast/Orbit^ to identify cells overexpressing CLASP^Mast/Orbit^, GFP-Rod signals could not be used to evaluate MT-kinetochore attachment status. Nevertheless, chromosome flipping frequency was significantly reduced in GFP-CLASP^Mast/Orbit^ overexpressing cells, supporting our hypothesis (Fig. 6C–E, Movie 6). However, spindle length, chromosome alignment, and mitotic duration were not restored by GFP-CLASP^Mast/Orbit^ overexpression (Fig. S4). The suppression was thus specific to MT attachment stability.

### GFP-Mad2 accumulation after KIF18A RNAi in HeLa cells

Several studies have characterised loss-of-function phenotypes for human kinesin-8, KIF18A (Janssen et al., 2018; Kim and Stumpff, 2018; Mayr et al., 2007; Stumpff et al., 2008; Stumpff et al., 2012). The components of the spindle assembly checkpoint (SAC) have been also analysed in a few studies using fixed cells (Janssen et al., 2018; Kim and Stumpff, 2018; Mayr et al., 2007). To confirm the phenotype and also possibly gain new insights into KIF18A function, we performed our own RNAi analysis using living HeLa cells expressing GFP-Mad2: Mad2 is a major component of SAC, and is rapidly recruited to the kinetochores that are not properly attached to MTs (Joglekar, 2016). For example, upon laser cutting of kinetochore MTs, Mad2 signals appeared within a few minutes (Dick and Gerlich, 2013).

We first confirmed that punctate signals of GFP-Mad2 were rarely detected on aligned chromosomes but were often observed in prometaphase, during which the majority of the kinetochores were unattached to MTs (Fig. 7A). They were also detectable when spindle MTs at metaphase were depolymerised with nocodazole (Fig. 7B).

**Figure 7.**
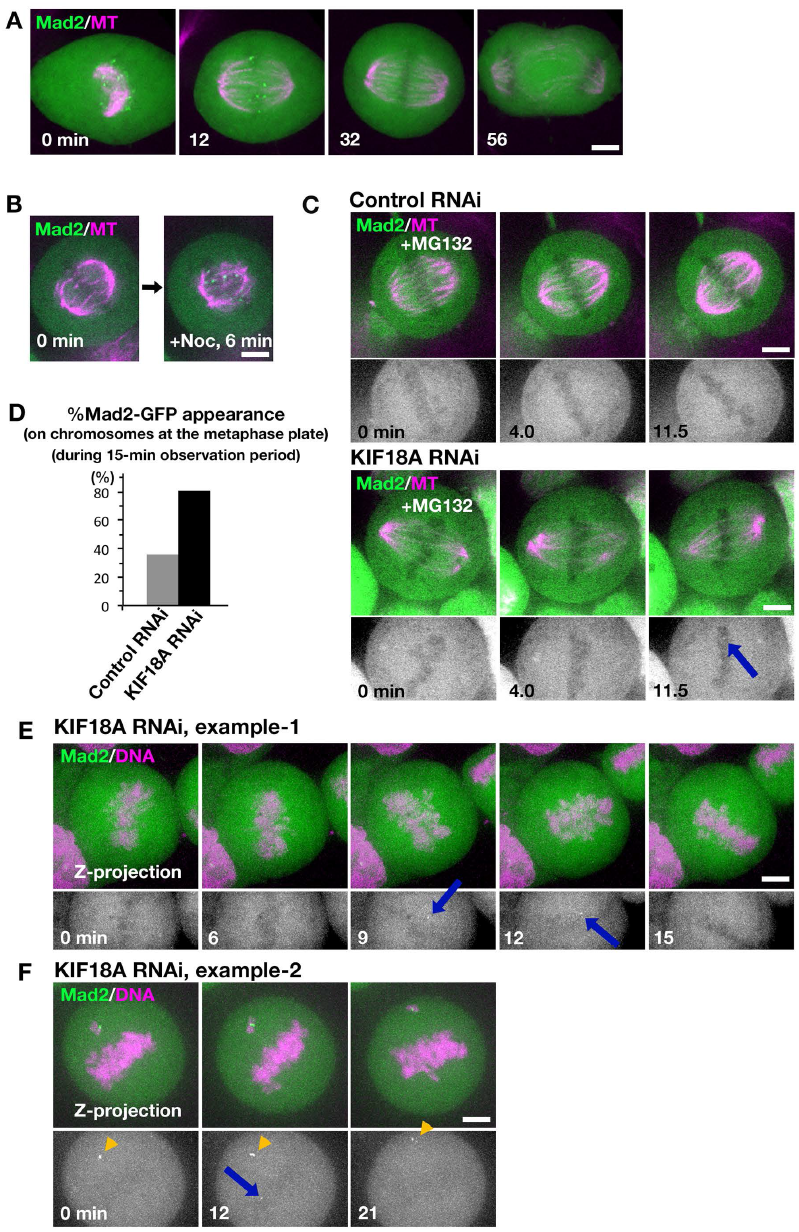
GFP-Mad2 dynamics in the absence of KIF18A in HeLa cells. **(A)** GFP-Mad2 localisation during mitosis in the control HeLa cell. Images were acquired from eleven z-sections (separated by 2 μm) and are displayed after maximum projection. **(B)** Accumulation of GFP-Mad2 on kinetochores upon MT depolymerisation by nocodazole. Image was acquired every 30 s at a single focal plane. **(C)** Metaphase-arrested cells were imaged every 30 s at a single focal plane after control or KIF18A RNAi. Arrow indicates the appearance of a punctate GFP-Mad2 signal at the metaphase plate. **(D)** Frequency of the cells in which GFP-Mad2 appeared on chromosomes at the metaphase plate during the 15-min observation period. N = 47 each. **(E, F)** GFP-Mad2 appearance at the metaphase plate in KIF18 RNAi-treated cells (arrows). As a reference, brighter GFP-Mad2 signals on persistently unaligned chromosomes are shown in F (arrowheads). Maximum projection images are shown. Bars, 5 μm.

Second, we observed GFP-Mad2 every 4 s in the monopolar spindle assembled by inhibiting kinesin-5 in the presence or absence of KIF18A, which was an assay analogous to those performed in S2 cells. However, in HeLa, GFP-Mad2 was constantly observed and chromosomes were dynamic even in the control cells: we could not identify a defect in the KIF18A RNAi-treated cells by our eyes (n = 41 cells; Movie 7).

We then observed GFP-Mad2 dynamics in the bipolar spindle in the absence of KIF18A in two conditions. First, we treated cells with MG132 to arrest them at metaphase and performed imaging of a single focal plane of multiple mitotic cells every 30 s for 15 min (Fig. 7C). We analysed a total of 47 metaphase cells for control and KIF18A-depeleted cells. In this artificially-arrested condition, unaligned chromosomes with strong punctate Mad2 signals were observed in control and KIF18A RNAi samples at similar frequencies (40% and 36%). In contrast, Mad2 signals also appeared on aligned chromosomes — often transiently — in 81% of the cells missing KIF18A, while only 36% of the control cells displayed such behaviour (Fig. 7C, D; arrow indicates the GFP-Mad2 signal). The results suggested that, in the absence of KIF18A, a subpopulation of seemingly aligned chromosomes had an improper attachment. Moreover, the transient signal appearance suggested that Mad2-negative kinetochores (i.e. properly attached kinetochores) occasionally altered their attachment status and became Mad2-positive. Alternatively, the Mad2 appearance/disappearance might simply reflect the kinetochore motility in and out of focal plane of the microscope.

To further test if Mad2-negative chromosomes could become Mad2 positive in the absence of KIF18A, we acquired z-stack images of normally cycling cells every 3 min for 30 min after RNAi (0.5 µm separation, >30 sections, by which the entire chromosome sets were covered at the beginning of imaging). Interestingly, in 70% cells (n = 23), GFP-Mad2 signals appeared on chromosomes which had not displayed any signals in previous time frames (Fig. 7E, F, blue arrows). The signal intensity was generally weaker than that observed on persistently unaligned chromosomes (yellow arrowheads in Fig. 7F), supporting the idea that the GFP-Mad2 emerged on those chromosomes during metaphase instead of persisting from early prometaphase. From these results, we concluded that once-disappeared GFP-Mad2 could reemerge in the absence of KIF18A. Thus, KIF18A may be required to maintain proper attachment status between MTs and kinetochores.

## Discussion

### Biochemical activity of kinesin-8^Klp67A^

We observed the processive plus-end-directed motility, catastrophe-induction, and rescue/pausing activities of kinesin-8^Klp67A^ towards MTs. This combination of activities has not been observed for another five *Drosophila* MT plus-end-regulating proteins we have characterised so far using an identical assay (Li et al., 2012; Moriwaki and Goshima, 2016). Motility and catastrophe induction can be deduced from the amino acid sequences of Klp67A’s motor domain; the motor has been shown to be critical in a previous study using a rigor mutant (Savoian and Glover, 2010). In contrast, how rescue/pausing activity and also mild growth-accelerating activity are executed remains unclear; it might involve the usage of the tubulin-binding region next to the motor domain to increase the local concentration of tubulin.

Whether human KIF18A is a depolymerase (Locke et al., 2017; Mayr et al., 2007) or MT dynamics suppressor (Du et al., 2010) has been debated. However, those studies used distinct assay and buffer conditions *in vitro*; it is possible that KIF18A can execute both activities in cells, namely, being endowed with a similar set of activities to *Drosophila* Klp67A. Budding yeast Kip3 is an established MT depolymerase, but rescue and pausing frequencies are also reduced in the mutant *in vivo* (Gupta et al., 2006), for which the tubulin-binding region is involved (Su et al., 2011). Our study suggests that the multiple activities observed here for kinesin-8^Klp67A^ in a single experimental condition are widely conserved among kinetochore-localised kinesin-8s.

### Kinesin-8^Klp67A^ is required for stable kinetochore-MT attachment

Previous studies using RNAi or mutants of kinesin-8^Klp67A^ consistently reveal its role in spindle length regulation (Buster et al., 2007; Gatt et al., 2005; Goshima and Vale, 2003; Goshima and Vale, 2005; Goshima et al., 2005; Savoian et al., 2004; Wang et al., 2010). This was confirmed in this study, and most likely is attributed to the catastrophe-inducing function. However, experiments involving colcemid treatment indicated that MT elongation with reduced catastrophe is not the causal factor leading to chromosome misalignment. GFP-Rod imaging further uncovered a specific role of kinesin-8^Klp67A^ at the kinetochore-MT interface: it ensures persistent MT attachment to the kinetochore.

Why are MTs frequently detached in the absence of kinesin-8^Klp67A^? We observed that overexpression of the MT rescue/pausing factor CLASP largely rescued the flipping behaviour of chromosomes. Since flipping is associated with the detachment event, we interpret that CLASP overexpression reduced MT detachment rates. The data therefore suggest that kinetochores are prone to detach from MTs during rapid and persistent depolymerisation. On the other hand, *in vitro* study using Ndc80-decorated beads and depolymerising MTs indicated robust load-bearing attachment of the beads during depolymerisation (Powers et al., 2009). However, this observation might be reconciled with our findings, as Aurora B kinase could dampen Ndc80’s MT binding ability in cells. Consistent with this notion, we observed stable MT association when Aurora B kinase was inhibited. We propose that kinesin-8^Klp67A^ constitutes an additional layer of the MT attachment interface.

### Functional similarity and difference between *Drosophila* and human kinesin-8s

It was previously reported that KIF18A depletion increases the amplitude of chromosome oscillation, where MT dynamics regulation, rather than attachment per se, is defective (Stumpff et al., 2008). This behaviour is consistent with the *in vitro* MT stabilisation activity (Du et al., 2010). However, a more recent study suggested that kinetochore-MT interaction is also perturbed in this condition (Kim and Stumpff, 2018). The study attributes the phenotype partly to defects in KIF18A-dependent PP1 delocalisation (Kim and Stumpff, 2018); however, those investigations also showed the presence of a PP1-independent function of KIF18A for chromosome alignment. It remains to be determined if PP1 mis-localisation contributes to chromosome misalignment in the absence of kinesin-8^Klp67A^, which does not possess the PP1-binding motif (De Wever et al., 2014).

While we were revising the manuscript, a new report on human KIF18A function was published, in which KIF18A depletion phenotype was investigated in a haploid cell line as well as HeLa cells (Janssen et al., 2018). In this report, MTs and Mad1 (binding partner of Mad2) were immuno-stained in the metaphase-arrested cell in the absence of KIF18A and three interesting conclusions have been drawn: 1) Mad1 is detected on multiple, but not all, kinetochores in the absence of KIF18A, which was consistent with a previous report using Mad2 (Mayr et al., 2007); 2) similar numbers of MTs are associated with each kinetochore, regardless of the presence or absence of Mad1 accumulation (i.e. MT-kinetochore attachment is not disrupted in the absence of KIF18A); 3) nevertheless, tension is not sufficiently applied to Mad1-positive kinetochores. Our data obtained in live cells confirmed the first point. We detected even fewer numbers of Mad2 signals than Janssen et al (2018); this might be the due to the difference in knockdown efficiency or methodology (live cell imaging vs. immunostaining after pre-extraction of the cell). In contrast, neither study has revealed the mechanism by which tension is reduced despite the apparently similar numbers of MTs attached in an end-on fashion to the kinetochore. However, it would be reasonable to assume that kinetochore-MT attachment is somewhat skewed in the absence of KIF18A. Our data suggests that this could occur even when attachment has been once established. In this regard, it is intriguing that we could not identify an apparent defect in the monopolar spindle. KIF18A may be particularly critical in the bipolar spindle, in which it might fine-tune the attachment mode for the kinetochore to generate sufficient tension.

Our data and those in Janssen et al. (2018) both agreed that MT attachment does not require KIF18A, which is consistent with the conclusion drawn from the analysis using *Drosophila* Klp67A. However, in the case of *Drosophila*, the loss of Klp67A caused MT detachment from the kinetochore, whereas the KIF18A depletion phenotype was milder. This was possibly due to the presence of other regulators of kinetochore-MT attachment in mammals, such as Ska and SKAP/astrin complexes, which are missing in *Drosophila* (Kern et al., 2017; Schmidt et al., 2012). Nevertheless, proper attachment does require KIF18A function, as reflected by the appearance of Mad1/Mad2 signals and the loss of tension in the absence of KIF18A (Janssen et al., 2018). Thus, KIF18A ensures the attachment mode for the kinetochore to generate sufficient tension. Several studies suggest an involvement of KIF18A in carcinogenesis (Hitti et al., 2016; Zhang et al., 2010). It would be intriguing to investigate how KIF18A contributes to chromosome segregation in various cancer cell models.

## Materials and methods

### RNAi and cell line selection

S2 cell culture and RNAi were performed as previously described (Bettencourt-Dias and Goshima, 2009; Goshima et al., 2007; Ito and Goshima, 2015). In brief, Schneider’s medium (Gibco) supplemented with 10% serum was used for cell culture. Cell lines were selected with hygromycin or puromycin following plasmid transformation with the TransIT-Insect reagent (Takara). Plasmids used in this study are listed in Table S1, whereas dsRNA sequences employed here are available at (Goshima and Vale, 2005; Goshima et al., 2007). For RNAi experiments, cells were treated with dsRNAs for 3–4 days and then plated on concanavalinA-coated glass-bottom dishes for microscopy. HeLa cells stably expressing GFP-Mad2 (a gift of Dr. Tomomi Kiyomitsu [Nagoya University]) were cultured in DMEM with 10% serum. RNAi was conducted with RNAiMax (Invitrogen) and 25 nM siRNA previously described (UAAAUUACCCGAACAAGAAtt; (Tanenbaum et al., 2009)). Imaging was started at 24 h.

### Microscopy

S2 MTs were stained with 15 nM SiR-tubulin, whereas EB1-SNAP was visualized with 15–30 nM SiR-SNAP. Live S2 imaging was performed with a spinning-disc confocal microscope (Nikon Ti; 100× 1.45 NA or 60 × 1.40 NA lens, EMCCD camera ImagEM [Hamamatsu], CSU-X1 [Yokogawa]). A TIRF microscope was used in the *in vitro* MT dynamics experiment (Nikon Ti; 100× 1.49 NA lens, EMCCD camera Evolve [Roper]). 488/561/640-nm excitation lasers were associated with both microscopes. Microscopes were controlled by micromanager and images were processed with ImageJ. For the colcemid treatment experiment in Fig. 5D, S2 cells expressing GFP-Rod and H2B-mCherry were stained with 15 nM SiR-tubulin and treated with 25 µM MG132 (≥ 30 min). Cells were imaged every 1 min for 20 min, and 5 µg/mL colcemid was supplied at 2 min. In Fig. S3B–D, cells were treated with 60 ng/mL colcemid for 2 h; images were acquired every 2 min. All imaging was performed at approximately 25°C. HeLa cells expressing GFP-Mad2 by CMV promoter were stained with 50 nM SiR-tubulin. Metaphase arrest, MT depolymerisation, and monopolar spindle formation were induced by 25 µM MG132, 100 nM nocodazole, and 5 µM STLC (a kinesin-5 inhibitor), respectively (≥ 1 h).

### Protein purification

S2 tubulin was purified by using a previously described method, using GST-tagged TOG1 domain (*S. cerevisiae* Stu2, 1–306 a.a.) (Moriwaki and Goshima, 2016; Widlund et al., 2012). Klp67A-GFP-6His was expressed in *E. coli* SoluBL21 cells (250 mL culture in L-rich medium, 0.1–0.5 mM IPTG at 18°C for 16–20 h). Cells were resuspended in Lysis Buffer (50 mM MOPS-NaOH [pH = 7.2], 250 mM NaCl, 2 mM MgCl_2,_ 1 mM EGTA, 0.5 mM Phenylmethylsulfonyl fluoride (PMSF), peptide cocktail (1 µg/ml leupeptin, pepstatin, chymostatin, and aprotinin), 2 mM 2-mercaptoethanol, 0.1 mM ATP), sonicated with a homogeniser (Branson, 450DA), bound to Ni-NTA (4°C, 60–90 min), washed with Wash Buffer (Lysis Buffer supplemented with 20 mM imidazole and 0.2 % Tween), followed by elution 5–8 times with Elution Buffer (MRB80 [80 mM PIPES-KOH (pH = 6.8), 1 mM EGTA, 4 mM MgCl_2_], 100 mM KCl, 250 mM imidazole, 2 mM 2-mercaptoethanol, 1 mM ATP). The eluate was subjected to sucrose gradient sedimentation. Gel filtration chromatography was not used because the procedure caused protein loss due to unknown reasons. A 2.5- 40 % sucrose gradient was made with 2 mL buffer (MRB80, 100 mM KCl, 0.1 mM ATP, 1mM DTT, sucrose) in a 2.2 mL, 11 × 35 mm ultracentrifugation tube (Beckman Coulter, # 347357). Protein solution (200 µL) was applied and centrifuged with a TLS-55 rotor (214,000 × g, 4 h, 4°C), and 16 fractions were collected. Fractions containing Klp67A-GFP-6His were identified with SDS-PAGE and Coomassie staining, followed by flash freezing. The tail-less Klp67A [1–612 a.a.]-GFP-6His and motor-alone Klp67A [1–359 a.a.]-GFP-6His were purified in a manner identical to full-length Klp67A-GFP-6His, except that the gradient sedimentation step was omitted and the buffer was exchanged instead to MRB80 containing 100 mM KCl, 20 % sucrose, 0.1 mM ATP, and 1 mM DTT, using the desalting column PD MiniTrap G-25 (GE Healthcare). The solution was flash frozen with liquid nitrogen and stored at -80 °C. 6His-Klp10A was purified with Ni-NTA, as previously described (Moriwaki and Goshima, 2016).

### Sucrose gradient sedimentation of the Klp67A-tubulin complex

Purified Klp67A-GFP-6His (~2 µM) solution was dialysed with Tube-O-DIALYZE, Micro, 8K MWCO (TaKaRa) in MRB80 containing 75 mM KCl, 5 % sucrose, 0.1 mM ATP, 1mM DTT for 6 h at 4 °C. The dialysed Klp67A was mixed with 4 µM pig tubulin, 1 mM ATP, and 1mM GTP, and incubated for 10 min at room temperature. The mixed solution was loaded into sucrose gradient buffer (MRB80, 75 mM KCl, 1 mM ATP, 1mM GTP, 1mM DTT, sucrose) to make 5–30 % sucrose gradient in a 2.2 mL, 11 × 35 mm ultracentrifugation tube and centrifuged with TLS-55 rotor (214,000 × *g*, 5.5 h, 4°C). The sedimented solution was divided into 16 fractions and the peak fractions were identified with SDS-PAGE and Sypro Ruby Staining.

### In vitro MT polymerisation assay

The *in vitro* MT polymerisation assay was performed essentially following a method previously described (Li et al., 2012; Moriwaki and Goshima, 2016). A silanized coverslip was coated with anti-biotin (1–5% in 1 × MRB80, Invitrogen), and the nonspecific surface was blocked with Pluronic F127 (1% in 1 × MRB80, Invitrogen). Biotinylated MT seeds (50–100 µM tubulin mix containing 10% biotinylated pig tubulin and 10% Alexa647-labelled pig tubulin with 1 mM GMPCPP) were specifically attached to the functionalised surface by biotinylated tubulin-anti-biotin links. After the chamber was washed with 1 × MRB80, MT growth was initiated by flowing 10 µM tubulin (containing 80% S2 tubulin and 20% Alexa568-labelled pig tubulin) and Klp67A-GFP into the assay buffer (1 × MRB80, 75 mM KCl, 1 mM GTP, 1 mM ATP, 0.5 mg/mL κ-casein and 0.1% methylcellulose, 5.5 % sucrose), and an oxygen scavenger system (50 mM glucose, 400 µg/mL glucose-oxidase, 200 µg/mL catalase, and 4 mM DTT). The samples were sealed with candle wax. During experiments, the samples were maintained at approximately 25°C, and images were collected every 3 s for 15 min using TIRF microscopy. MT grew from both ends, but only the plus-end dynamics were analysed. MT depolymerisation assay was conducted in an identical condition, except that no tubulin was included. MT gliding assay was performed following a previous report (Miki et al., 2015) with slight modification to the buffer. Briefly, the flow chamber was washed with 1 × MRB80 and purified kinesin-1 motor (K560-6His) was flowed into the chamber. After washing with MRB80 containing 0.5 mg/ml k-casein, the motility buffer (1× MRB80, 75 mM KCl, GMPCPP-stabilized MTs with Alexa 647-labels, 1 mM ATP, 0.5 mg/ml k-casein and 0.1% methylcellulose, 5% sucrose), with an oxygen scavenger system [50 mM glucose, 400 mg/ml glucose oxidase, 200 mg/ml catalase and 4 mM dithiothreitol (DTT)] and 4 nM Klp67A-GFP, was flowed into the chamber. The single kinesin motility assay was conducted following other publications (Naito and Goshima, 2015) with slight modification to the buffer. A silanized coverslip was coated with anti-biotin antibody, and a solution containing 1% pluronic acid was loaded into the chamber. After washing once with 1 × MRB80, GMPCPP-stabilised MTs labelled with Alexa 647 and biotin were loaded. After a 1 × MRB80 wash, Klp67A-GFP was loaded into the chamber with a buffer identical to that used for the *in vitro* MT polymerisation assay. GFP was bleached for 3 min with 20 mW, 488 nm laser (maximum power), followed by imaging unbleached GFP with 40% power.

### Data analysis

MT attachment instability was determined by counting the detachment events over time (monotelic to unattachment/lateral interaction or syntelic to monotelic conversion), whereas MT attachment frequency was obtained for monotelically-attached chromosomes. MT plus-end dynamics *in vitro* were analysed based on kymographs, following others (Moriwaki and Goshima, 2016): catastrophe frequency was determined by dividing the number of shrinkage events by the sum of growth and pause times, whereas the transition from shrinkage to pause or growth was considered a rescue event and the rescue frequency (for shrinkage time) was calculated. When MTs did not grow or shrink more than two pixels (0.32 µm) for five or more frames (15 s), this period was defined as a pause. Intensity of fluorescent tubulin on GMPCPP-MT was measured with the line tool associated with Fiji/ImageJ (images acquired at 5 min of time-lapse were used). The obtained value was divided by the fluorescent intensity of GMPCPP-MT. To investigate the correlation between Klp67A-GFP intensity at the tip and the growth/shrinkage rate, kymographs were analysed with the segmented line tool associated with Fiji/ImageJ. At each time point, the maximum GFP intensity within 5 pixels (0.8 µm) around the tip was obtained and, after background intensity subtraction, the mean GFP intensity during the growth (or shrinkage) was calculated. The plot was analysed with general linear regression, where GFP intensity and growth/shrinkage rate were used as explanatory and response variables respectively, and gamma distribution was assumed. Reciprocal function was used as link function. To make the response variable exclusively positive values, 0 µm/min growth rate was approximated to 1 × 10^-9^ µm/min. The approximate curve was drawn with 95% Wald confidence interval. P values were obtained with likelihood-ratio test. Image analysis was performed with Jython scripts associated with Fiji/ImageJ. Further statistical analysis and data visualizations were performed on R or Prism.

## Acknowledgements

We thank Kosuke Ariga for helping data analysis, Tomoko Nishiyama for technical support, Tomomi Kiyomitsu for valuable comments on the manuscript, and Elsa Tungadi for proofreading. This work was funded by JSPS KAKENHI (15KT0077, 17H01431) and Laura and Arthur Colwin Endowed Summer Research Fellowship Fund (2015) of the Marine Biological Laboratory to G.G. T.E. is a recipient of a JSPS pre-doctoral fellowship (16J02807).

## Author contributions

T.E. and G.G. conceived and designed the research project. T.E. performed most of the experiments and analysed the data. G.G. performed some experiments, analysed the data, and wrote the paper.

## Supplemental Movie Legends

**Movie 1. Processive motility and plus-end-accumulation of kinesin-8^Klp67A^**

Kinesin-8^Klp67A^-GFP was imaged every 1 s with TIRF microscopy. We first illuminated the imaging field for ~3 min with high laser power to bleach most, if not all, kinesin-8^Klp67A^-GFP at the microtubule end, and then restarted time-lapse imaging with a normal laser power. Bar, 5 µm.

**Movie 2. MT attachment stability in the monopolar spindle in the presence or absence of kinesin-8^Klp67A^**

GFP-Rod (green), H2B-mCherry (blue), and SiR-tubulin (magenta) images were acquired every 3 s with spinning-disc confocal microscopy in kinesin-5^Klp61F^ single or kinesin-8^Klp67A^/kinesin-5^Klp61F^ double RNAi-treated cells. Bar, 5 µm.

**Movie 3. MT attachment stability in the bipolar spindle in the presence or absence of kinesin-8^Klp67A^**

GFP-Rod (green), H2B-mCherry (blue), and SiR-tubulin (magenta) images were acquired every 3 s with spinning-disc confocal microscopy in kinesin-8^Klp67A^ or control RNAi-treated cells. Bar, 5 µm.

**Movie 4. MT attachment stability in the absence of augmin^Dgt6^**

GFP-Rod (green), H2B-mCherry (blue), and SiR-tubulin (magenta) images were acquired every 3 s with spinning-disc confocal microscopy in augmin^Dgt6^/kinesin-5^Klp61F^ RNAi-treated cells. Bar, 5 µm.

**Movie 5. Stable syntelic attachment in the monopolar spindle after Aurora B inhibition in the absence of kinesin-8^Klp67A^**

GFP-Rod (green), H2B-mCherry (blue), and SiR-tubulin (magenta) images were acquired every 3 s with spinning-disc confocal microscopy in augmin^Dgt6^/kinesin-5^Klp61F^ RNAi-treated cells. Twenty µM Binucleine 2 (*Drosophila* Aurora B inhibitor) or control DMSO was added at time 0. Bar, 5 µm.

**Movie 6. CLASP^Mast/Orbit^ overexpression rescued MT attachment stability in the absence of kinesin-8^Klp67A^**

GFP-Rod (green), GFP-CLASP^Mast/Orbit^ (green), H2B-mCherry (blue), and SiR-tubulin (magenta) images were acquired every 3 s with spinning-disc confocal microscopy in kinesin-8^Klp67A^/kinesin-5^Klp61F^ double RNAi-treated cells expressing GFP-CLASP^Mast/Orbit^. Bar, 5 µm.

**Movie 7. GFP-Mad2 dynamics in the monopolar spindle after KIF18A RNAi in HeLa cells**

GFP-Mad2 (green) and SiR-tubulin (magenta) images were acquired every 4 s with spinning-disc confocal microscopy in kinesin-8^KIF18A^ and control RNAi-treated cells that were treated with STLC, an inhibitor of kinesin-5^Eg5^. Nine randomly selected cells are displayed for each sample. Areas devoid of fluorescence represent chromosomes. Neither of the authors could identify any phenotype associated with kinesin-8^KIF18A^ depletion. Bar, 5 µm.

**Table S1.** Plasmids used in this study Name Insert.

| Name | Insert | Vector | Note |
| --- | --- | --- | --- |
| pED128 | Klp67A[full-length]-GFP-6His | pET23 |  |
| pED182 | Klp67A[1–612a.a.]-GFP-6His | pET23 |  |
| pED183 | Klp67A[1–359a.a.]-GFP-6His | pET23 |  |
| pGG952 | 6His-Klp10A | pDEST17 | Moriwaki and Goshima (2016) |
| pED158 | sfGFP-Rod | pAc |  |
| pGFP-CLASP | GFP-Mast/Orbit | pMT | Goshima et al. (2007) |
| pGG482 | H2B-mCherry | pAc |  |
| pED309 | EB1-SNAP | pAc |  |
| pCoHygromycin | HygR | Copia |  |

**Figure S1.**
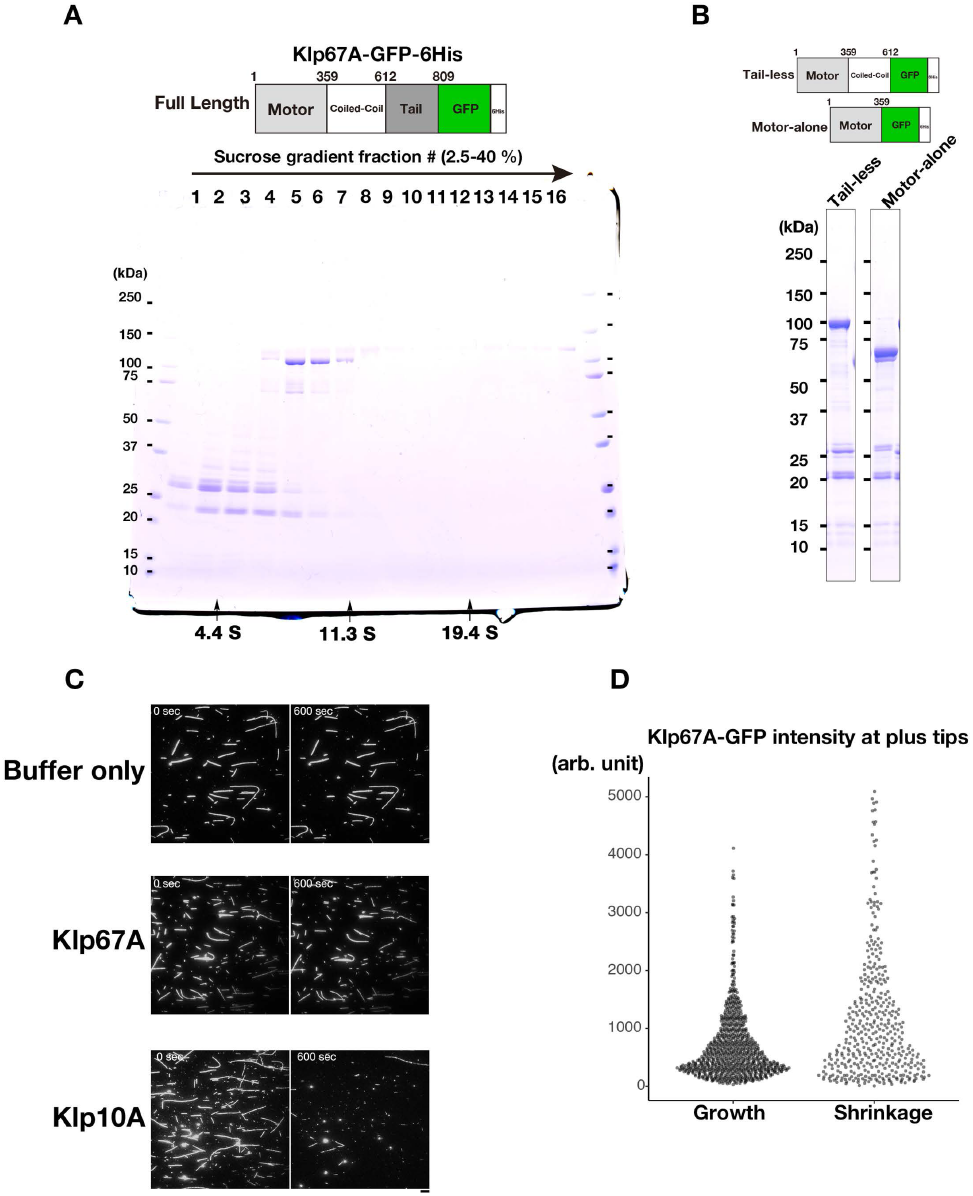
Purification of kinesin-8^Klp67A^-GFP. **(A)** Coomassie staining of full-length kinesin-8^Klp67A^ tagged with GFP and 6 x His after sucrose gradient centrifugation. BSA, catalase, and thyroglobulin were used as the markers (indicated by arrows). Fraction 6 was used for activity measurement. **(B)** Coomassie staining of purified, truncated kinesin-8^Klp67A^ tagged with GFP and 6 x His. **(C)** Kinesin-8^Klp67A^ (10 nM) cannot depolymerise GMPCPP-stabilised MTs. MTs stabilised with GMPCPP were mixed with full-length kinesin-8^Klp67A^ (10 nM) and, as a positive control, MT depolymerase kinesin-13^Klp10A^ (10 nM). Bar, 5 μm. **(D)** Kinesin-8^Klp67A^-GFP intensities at the plus ends of growing and shrinking MTs (n = 891 and 409). The intensity was, on average, slightly higher on shrinking MTs (p < 0.001, Brunner-Munzel test).

**Figure S2.**
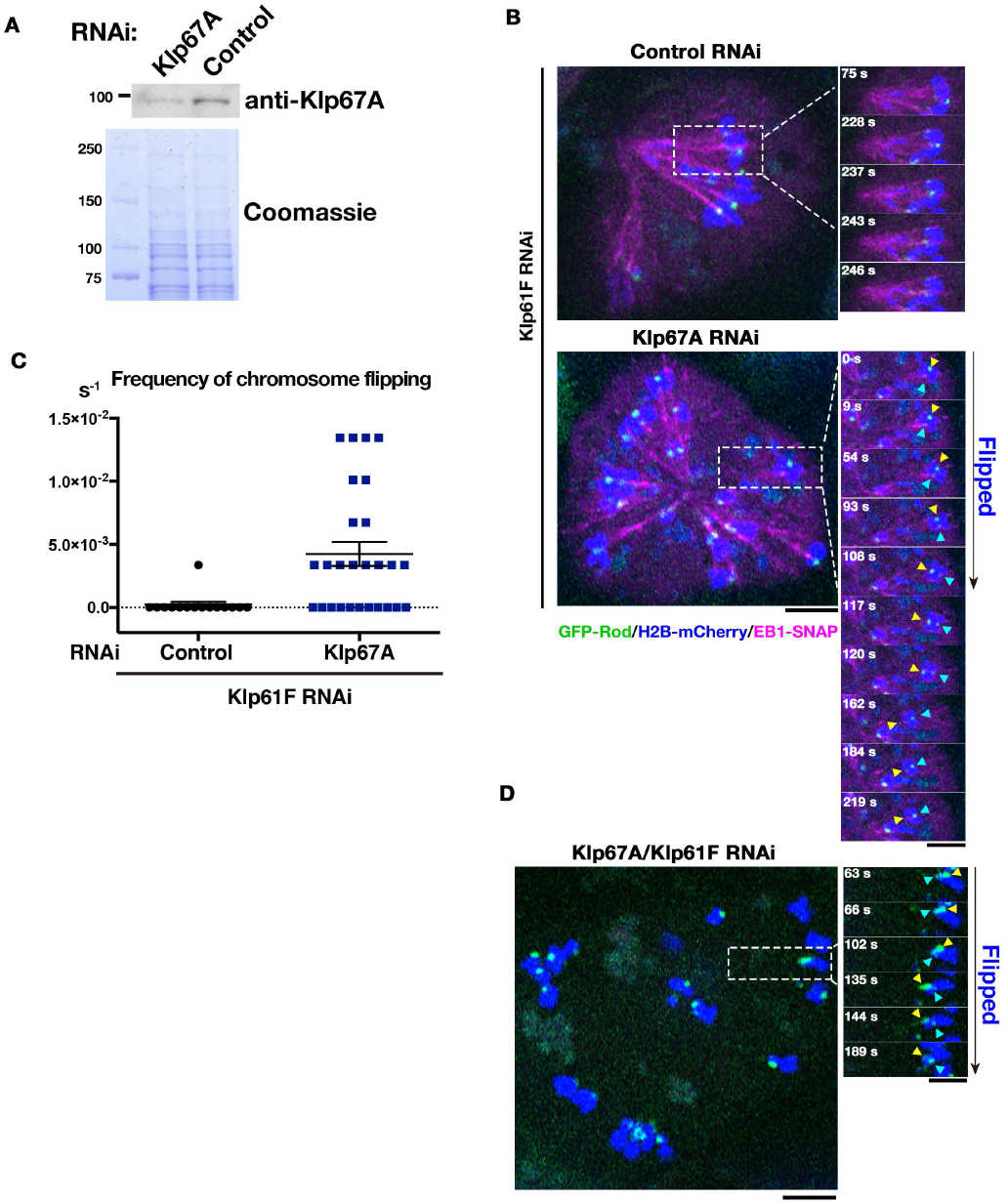
Kinesin-8^Klp67A^ RNAi causes MT attachment instability. **(A)** Immunoblotting confirmed 80% reduction of kinesin-8^Klp67A^ protein after RNAi (GFP-CLASP^Mast/Orbit^ expressing line was used). **(B-D)** MT attachment instability was not an artefact of SiR-tubulin staining, as the phenotype was reproduced without SiR-tubulin staining. In **(B)**, MTs were visualized by expressing fluorescently labeled EB1, whereas MTs were not labelled in **(D)**. **(C)** Chromosome flipping frequency was quantified (per cell, per sec, ±SEM). 15 cells and 27 cells were analysed (control and Klp67A RNAi). Bars, 5 μm.

**Figure S3.**
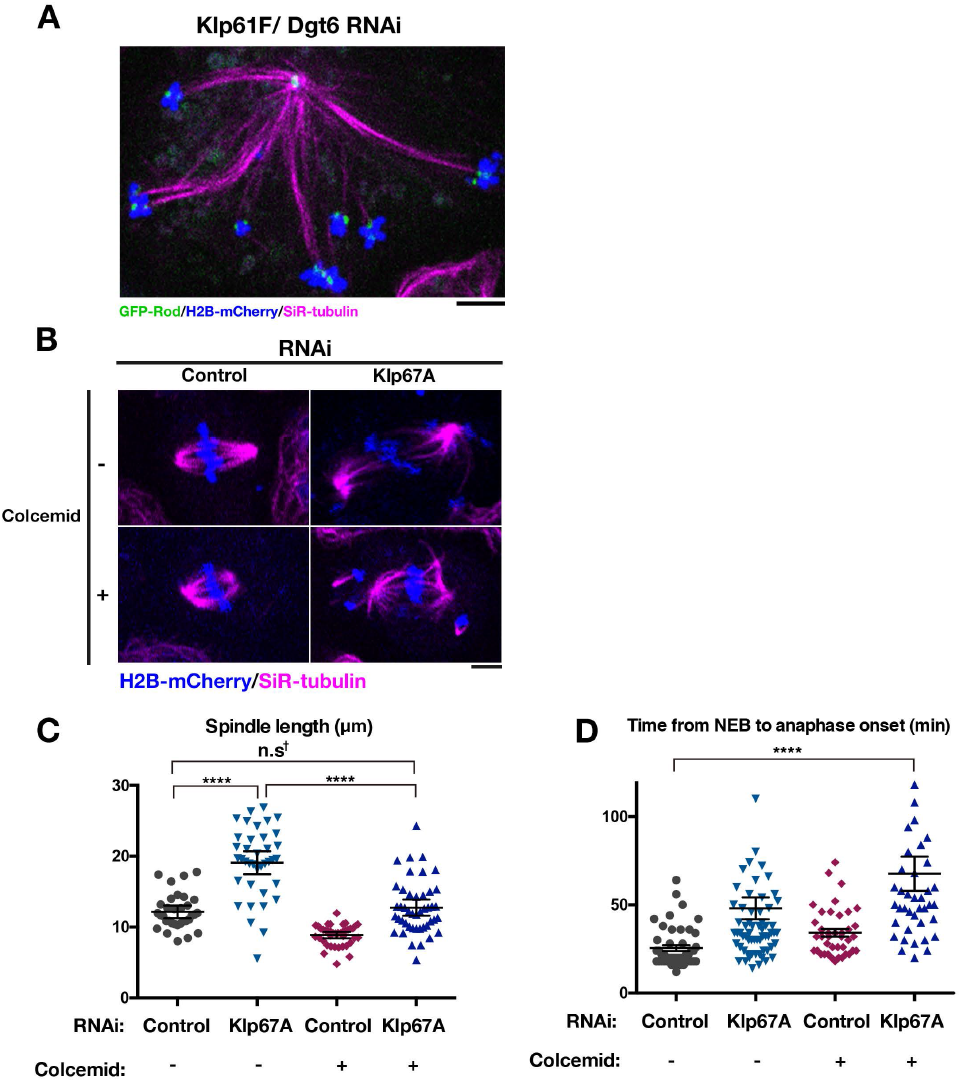
Artificial MT shortening cannot rescue chromosome misalignment induced by kinesin-8^Klp67A^ depletion. **(A)** Elongated monopolar spindle after double RNAi knockdown of augrnin^Dgt6^ and kinesin-5^Klp61F^. GFP-Rod signals indicated the presence of both monotelic and syntelic chromosomes. See Movie 4 for chromosome/kinetochore dynamics. **(B-D)** Force-shortening of the spindle by colcemid treatment (60 ng/mL) in kinesin-8^Klp67A^ RNAi-treated cells did not recover chromosome misalignment **(B)** and mitotic progression **(D)**, despite that the spindle was shortened to the control level **(C)**. Spindle length was measured at 16 min after NEBD. Marked with † is the comparison between control RNAi and Klp67A RNAi + colcemid: mean difference = 0.59, 95 % confidence interval on the difference = [-1.28, 2.46], **** indicates significant (p < 0.001) difference by Games-Howell test. N = 33, 38, 45 and 40 (C; from left to right), and 46, 66, 40, 41 (D, from left to right). Error bars indicate SEM. Experiments were performed twice, and the data from one experiment is presented. Two outliers obtained in kinesin-8^Klp67A^ RNAi-treated cells in **(D)** are not described in the graph but were taken into account during mean and SEM calculations. Bars, 5 μm.

**Figure S4.**
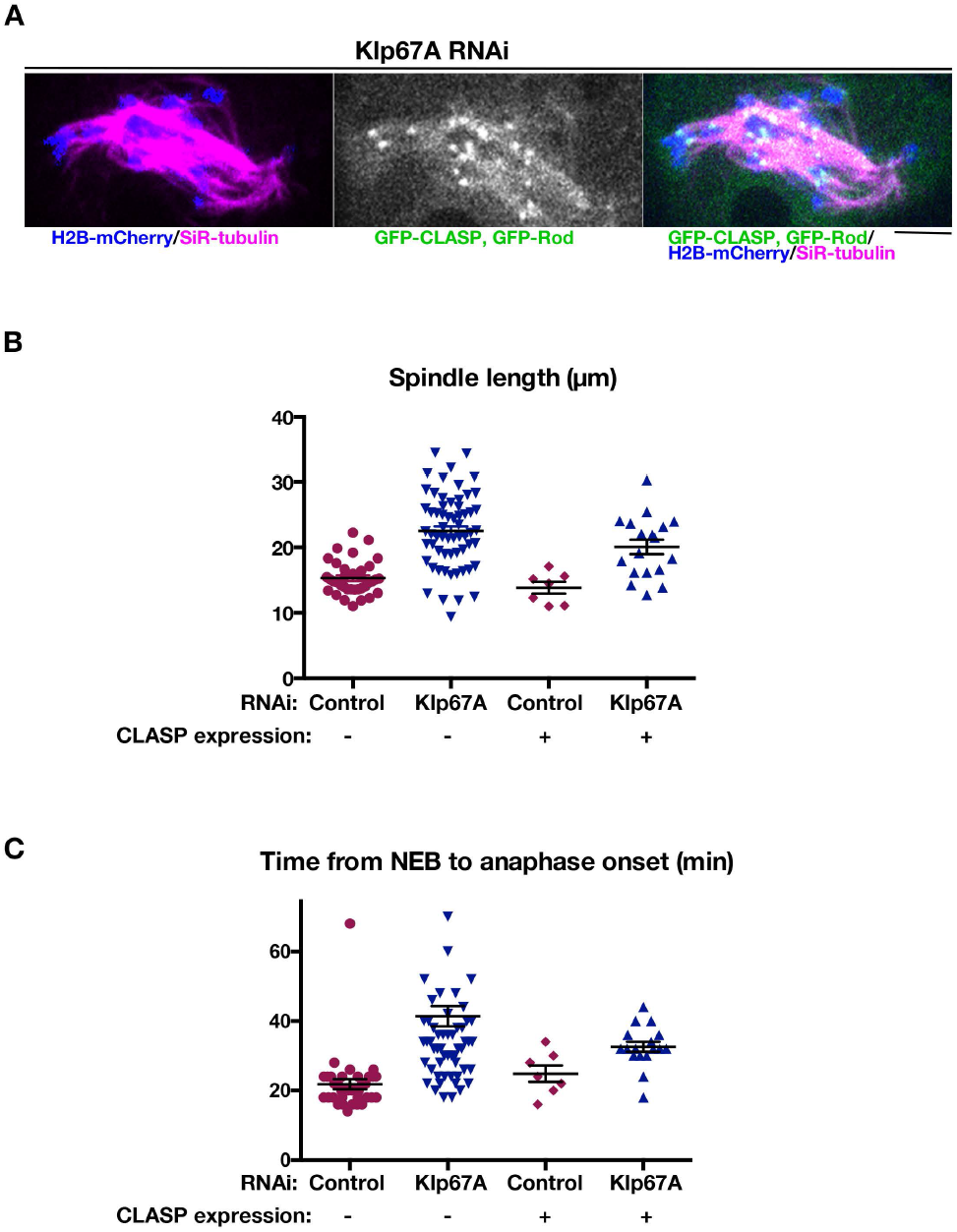
CLASP^Mast/Orbit^ overexpression cannot rescue chromosome misalignment induced by kinesin-8^Klp67A^ depletion. **(A)** A typical mitotic cell expressing GFP-CLASP^Mast/Orbit^ following kinesin-8^Klp67A^ RNAi. Chromosomes were severely misaligned in the spindle. **(B, C)** Metaphase spindle length and mitotic delay were also not rescued by CLASP^Mast/Orbit^ overexpression (error bars represents SEM).

## References

Al-Bassam, J., H. Kim, G. Brouhard, A. van Oijen, S.C. Harrison, and F. Chang. 2010. CLASP promotes microtubule rescue by recruiting tubulin dimers to the microtubule. Dev Cell. 19:245–258.

Basto, R., F. Scaerou, S. Mische, E. Wojcik, C. Lefebvre, R. Gomes, T. Hays, and R. Karess. 2004. In vivo dynamics of the rough deal checkpoint protein during Drosophila mitosis. Curr Biol. 14:56–61.

Bettencourt-Dias, M., and G. Goshima. 2009. RNAi in Drosophila S2 cells as a tool for studying cell cycle progression. Methods Mol Biol. 545:39–62.

Buster, D.W., D. Zhang, and D.J. Sharp. 2007. Poleward tubulin flux in spindles: regulation and function in mitotic cells. Mol Biol Cell. 18:3094–3104.

Cheeseman, I.M., S. Anderson, M. Jwa, E.M. Green, J. Kang, J.R. Yates, 3rd, C.S. Chan, D.G. Drubin, and G. Barnes. 2002. Phospho-regulation of kinetochore-microtubule attachments by the Aurora kinase Ipl1p. Cell. 111:163–172.

Cheeseman, I.M., J.S. Chappie, E.M. Wilson-Kubalek, and A. Desai. 2006. The conserved KMN network constitutes the core microtubule-binding site of the kinetochore. Cell. 127:983–997.

Cottingham, F.R., and M.A. Hoyt. 1997. Mitotic spindle positioning in Saccharomyces cerevisiae is accomplished by antagonistically acting microtubule motor proteins. J Cell Biol. 138:1041–1053.

De Wever, V., I. Nasa, D. Chamousset, D. Lloyd, M. Nimick, H. Xu, L. Trinkle-Mulcahy, and G.B. Moorhead. 2014. The human mitotic kinesin KIF18A binds protein phosphatase 1 (PP1) through a highly conserved docking motif. Biochem Biophys Res Commun. 453:432–437.

DeLuca, J.G., W.E. Gall, C. Ciferri, D. Cimini, A. Musacchio, and E.D. Salmon. 2006. Kinetochore microtubule dynamics and attachment stability are regulated by Hec1. Cell. 127:969–982.

Dick, A.E., and D.W. Gerlich. 2013. Kinetic framework of spindle assembly checkpoint signalling. Nat Cell Biol. 15:1370–1377.

Du, Y., C.A. English, and R. Ohi. 2010. The kinesin-8 Kif18A dampens microtubule plus-end dynamics. Curr Biol. 20:374–380.

Erent, M., D.R. Drummond, and R.A. Cross. 2012. S. pombe kinesins-8 promote both nucleation and catastrophe of microtubules. PLoS One. 7:e30738.

Gandhi, R., S. Bonaccorsi, D. Wentworth, S. Doxsey, M. Gatti, and A. Pereira. 2004. The Drosophila kinesin-like protein KLP67A is essential for mitotic and male meiotic spindle assembly. Mol Biol Cell. 15:121–131.

Garcia, M.A., L. Vardy, N. Koonrugsa, and T. Toda. 2001. Fission yeast ch-TOG/XMAP215 homologue Alp14 connects mitotic spindles with the kinetochore and is a component of the Mad2-dependent spindle checkpoint. EMBO J. 20:3389–3401.

Gatt, M.K., M.S. Savoian, M.G. Riparbelli, C. Massarelli, G. Callaini, and D.M. Glover. 2005. Klp67A destabilises pre-anaphase microtubules but subsequently is required to stabilise the central spindle. J Cell Sci. 118:2671–2682.

Gluszek, A.A., C.F. Cullen, W. Li, R.A. Battaglia, S.J. Radford, M.F. Costa, K.S. McKim, G. Goshima, and H. Ohkura. 2015. The microtubule catastrophe promoter Sentin delays stable kinetochore-microtubule attachment in oocytes. J Cell Biol. 211:1113–1120.

Goshima, G., M. Mayer, N. Zhang, N. Stuurman, and R.D. Vale. 2008. Augmin: a protein complex required for centrosome-independent microtubule generation within the spindle. J Cell Biol. 181:421–429.

Goshima, G., and R.D. Vale. 2003. The roles of microtubule-based motor proteins in mitosis: comprehensive RNAi analysis in the Drosophila S2 cell line. J Cell Biol. 162:1003–1016.

Goshima, G., and R.D. Vale. 2005. Cell cycle-dependent dynamics and regulation of mitotic kinesins in Drosophila S2 cells. Mol Biol Cell. 16:3896–3907.

Goshima, G., R. Wollman, N. Goodwin, J.M. Zhang, J.M. Scholey, R.D. Vale, and N. Stuurman. 2007. Genes required for mitotic spindle assembly in Drosophila S2 cells. Science. 316:417–421.

Goshima, G., R. Wollman, N. Stuurman, J.M. Scholey, and R.D. Vale. 2005. Length control of the metaphase spindle. Curr Biol. 15:1979–1988.

Gupta, M.L., Jr., P. Carvalho, D.M. Roof, and D. Pellman. 2006. Plus end-specific depolymerase activity of Kip3, a kinesin-8 protein, explains its role in positioning the yeast mitotic spindle. Nat Cell Biol. 8:913–923.

Hitti, E., T. Bakheet, N. Al-Souhibani, W. Moghrabi, S. Al-Yahya, M. Al-Ghamdi, M. Al-Saif, M.M. Shoukri, A. Lanczky, R. Grepin, B. Gyorffy, G. Pages, and K.S. Khabar. 2016. Systematic Analysis of AU-Rich Element Expression in Cancer Reveals Common Functional Clusters Regulated by Key RNA-Binding Proteins. Cancer Res. 76:4068–4080.

Ito, A., and G. Goshima. 2015. Microcephaly protein Asp focuses the minus ends of spindle microtubules at the pole and within the spindle. J Cell Biol. 211:999–1009.

Janssen, L.M.E., T.V. Averink, V.A. Blomen, T.R. Brummelkamp, R.H. Medema, and J.A. Raaijmakers. 2018. Loss of Kif18A Results in Spindle Assembly Checkpoint Activation at Microtubule-Attached Kinetochores. Curr Biol. 28:2685–2696 e2684.

Joglekar, A.P. 2016. A Cell Biological Perspective on Past, Present and Future Investigations of the Spindle Assembly Checkpoint. Biology (Basel). 5.

Kern, D.M., J.K. Monda, K.C. Su, E.M. Wilson-Kubalek, and I.M. Cheeseman. 2017. Astrin-SKAP complex reconstitution reveals its kinetochore interaction with microtubule-bound Ndc80. Elife. 6.

Kim, H., and J.K. Stumpff. 2018. Kif18A promotes Hec1 dephosphorylation to coordinate chromosome alignment with kinetochore microtubule attachment. bioRxiv. 304147.

Lampson, M.A., and E.L. Grishchuk. 2017. Mechanisms to Avoid and Correct Erroneous Kinetochore-Microtubule Attachments. Biology (Basel). 6.

Li, W., T. Moriwaki, T. Tani, T. Watanabe, K. Kaibuchi, and G. Goshima. 2012. Reconstitution of dynamic microtubules with Drosophila XMAP215, EB1, and Sentin. J Cell Biol. 199:849–862.

Locke, J., A.P. Joseph, A. Pena, M.M. Mockel, T.U. Mayer, M. Topf, and C.A. Moores. 2017. Structural basis of human kinesin-8 function and inhibition. Proc Natl Acad Sci U S A. 114:E9539–E9548.

Maiato, H., E.A. Fairley, C.L. Rieder, J.R. Swedlow, C.E. Sunkel, and W.C. Earnshaw. 2003. Human CLASP1 is an outer kinetochore component that regulates spindle microtubule dynamics. Cell. 113:891–904.

Maiato, H., A. Khodjakov, and C.L. Rieder. 2005. Drosophila CLASP is required for the incorporation of microtubule subunits into fluxing kinetochore fibres. Nat Cell Biol. 7:42–47.

Mayr, M.I., S. Hummer, J. Bormann, T. Gruner, S. Adio, G. Woehlke, and T.U. Mayer. 2007. The human kinesin Kif18A is a motile microtubule depolymerase essential for chromosome congression. Curr Biol. 17:488–498.

Mayr, M.I., M. Storch, J. Howard, and T.U. Mayer. 2011. A non-motor microtubule binding site is essential for the high processivity and mitotic function of kinesin-8 Kif18A. PLoS One. 6:e27471.

McHugh, T., A.A. Gluszek, and J.P.I. Welburn. 2018. Microtubule end tethering of a processive kinesin-8 motor Kif18b is required for spindle positioning. J Cell Biol.

Miki, T., M. Nishina, and G. Goshima. 2015. RNAi screening identifies the armadillo repeat-containing kinesins responsible for microtubule-dependent nuclear positioning in Physcomitrella patens. Plant Cell Physiol. 56:737–749.

Moriwaki, T., and G. Goshima. 2016. Five factors can reconstitute all three phases of microtubule polymerization dynamics. J Cell Biol. 215:357–368.

Musacchio, A., and A. Desai. 2017. A Molecular View of Kinetochore Assembly and Function. Biology (Basel). 6.

Naito, H., and G. Goshima. 2015. NACK kinesin is required for metaphase chromosome alignment and cytokinesis in the moss Physcomitrella patens. Cell Struct Funct. 40:31–41.

Powers, A.F., A.D. Franck, D.R. Gestaut, J. Cooper, B. Gracyzk, R.R. Wei, L. Wordeman, T.N. Davis, and C.L. Asbury. 2009. The Ndc80 kinetochore complex forms load-bearing attachments to dynamic microtubule tips via biased diffusion. Cell. 136:865–875.

Rogers, G.C., S.L. Rogers, T.A. Schwimmer, S.C. Ems-McClung, C.E. Walczak, R.D. Vale, J.M. Scholey, and D.J. Sharp. 2004. Two mitotic kinesins cooperate to drive sister chromatid separation during anaphase. Nature. 427:364–370.

Savoian, M.S., M.K. Gatt, M.G. Riparbelli, G. Callaini, and D.M. Glover. 2004. Drosophila Klp67A is required for proper chromosome congression and segregation during meiosis I. J Cell Sci. 117:3669–3677.

Savoian, M.S., and D.M. Glover. 2010. Drosophila Klp67A binds prophase kinetochores to subsequently regulate congression and spindle length. J Cell Sci. 123:767–776.

Schmidt, J.C., H. Arthanari, A. Boeszoermenyi, N.M. Dashkevich, E.M. Wilson-Kubalek, N. Monnier, M. Markus, M. Oberer, R.A. Milligan, M. Bathe, G. Wagner, E.L. Grishchuk, and I.M. Cheeseman. 2012. The kinetochore-bound Ska1 complex tracks depolymerizing microtubules and binds to curved protofilaments. Dev Cell. 23:968–980.

Smurnyy, Y., A.V. Toms, G.R. Hickson, M.J. Eck, and U.S. Eggert. 2010. Binucleine 2, an isoform-specific inhibitor of Drosophila Aurora B kinase, provides insights into the mechanism of cytokinesis. ACS Chem Biol. 5:1015–1020.

Straight, A.F., J.W. Sedat, and A.W. Murray. 1998. Time-lapse microscopy reveals unique roles for kinesins during anaphase in budding yeast. J Cell Biol. 143:687–694.

Stumpff, J., Y. Du, C.A. English, Z. Maliga, M. Wagenbach, C.L. Asbury, L. Wordeman, and R. Ohi. 2011. A tethering mechanism controls the processivity and kinetochore-microtubule plus-end enrichment of the kinesin-8 Kif18A. Mol Cell. 43:764–775.

Stumpff, J., G. von Dassow, M. Wagenbach, C. Asbury, and L. Wordeman. 2008. The kinesin-8 motor Kif18A suppresses kinetochore movements to control mitotic chromosome alignment. Dev Cell. 14:252–262.

Stumpff, J., M. Wagenbach, A. Franck, C.L. Asbury, and L. Wordeman. 2012. Kif18A and chromokinesins confine centromere movements via microtubule growth suppression and spatial control of kinetochore tension. Dev Cell. 22:1017–1029.

Su, X., H. Arellano-Santoyo, D. Portran, J. Gaillard, M. Vantard, M. Thery, and D. Pellman. 2013. Microtubule-sliding activity of a kinesin-8 promotes spindle assembly and spindle-length control. Nat Cell Biol. 15:948–957.

Su, X., W. Qiu, M.L. Gupta, Jr., J.B. Pereira-Leal, S.L. Reck-Peterson, and D. Pellman. 2011. Mechanisms underlying the dual-mode regulation of microtubule dynamics by Kip3/kinesin-8. Mol Cell. 43:751–763.

Tanenbaum, M.E., L. Macurek, A. Janssen, E.F. Geers, M. Alvarez-Fernandez, and R.H. Medema. 2009. Kif15 cooperates with eg5 to promote bipolar spindle assembly. Curr Biol. 19:1703–1711.

Tien, J.F., N.T. Umbreit, D.R. Gestaut, A.D. Franck, J. Cooper, L. Wordeman, T. Gonen, C.L. Asbury, and T.N. Davis. 2010. Cooperation of the Dam1 and Ndc80 kinetochore complexes enhances microtubule coupling and is regulated by aurora B. J Cell Biol. 189:713–723.

Tytell, J.D., and P.K. Sorger. 2006. Analysis of kinesin motor function at budding yeast kinetochores. J Cell Biol. 172:861–874.

Varga, V., J. Helenius, K. Tanaka, A.A. Hyman, T.U. Tanaka, and J. Howard. 2006. Yeast kinesin-8 depolymerizes microtubules in a length-dependent manner. Nat Cell Biol. 8:957–962.

Varga, V., C. Leduc, V. Bormuth, S. Diez, and J. Howard. 2009. Kinesin-8 motors act cooperatively to mediate length-dependent microtubule depolymerization. Cell. 138:1174–1183.

Wang, H., I. Brust-Mascher, D. Cheerambathur, and J.M. Scholey. 2010. Coupling between microtubule sliding, plus-end growth and spindle length revealed by kinesin-8 depletion. Cytoskeleton (Hoboken). 67:715–728.

Wargacki, M.M., J.C. Tay, E.G. Muller, C.L. Asbury, and T.N. Davis. 2010. Kip3, the yeast kinesin-8, is required for clustering of kinetochores at metaphase. Cell Cycle. 9:2581–2588.

Weaver, L.N., S.C. Ems-McClung, J.R. Stout, C. LeBlanc, S.L. Shaw, M.K. Gardner, and C.E. Walczak. 2011. Kif18A uses a microtubule binding site in the tail for plus-end localization and spindle length regulation. Curr Biol. 21:1500–1506.

West, R.R., T. Malmstrom, and J.R. McIntosh. 2002. Kinesins klp5(+) and klp6(+) are required for normal chromosome movement in mitosis. J Cell Sci. 115:931–940.

Widlund, P.O., M. Podolski, S. Reber, J. Alper, M. Storch, A.A. Hyman, J. Howard, and D.N. Drechsel. 2012. One-step purification of assembly-competent tubulin from diverse eukaryotic sources. Mol Biol Cell. 23:4393–4401.

Yu, N., L. Signorile, S. Basu, S. Ottema, J.H. Lebbink, K. Leslie, I. Smal, D. Dekkers, J. Demmers, and N. Galjart. 2016. Isolation of functional tubulin dimers and of tubulin-associated proteins from mammalian cells. Curr Biol. 26:1728–1736.

Zhang, C., C. Zhu, H. Chen, L. Li, L. Guo, W. Jiang, and S.H. Lu. 2010. Kif18A is involved in human breast carcinogenesis. Carcinogenesis. 31:1676–1684.

